# The arrested state of processing bodies supports mRNA regulation in early development

**DOI:** 10.1101/2021.03.16.435709

**Authors:** M. Sankaranarayanan, Ryan J. Emenecker, Marcus Jahnel, Irmela R. E. A. Trussina, Matt Wayland, Simon Alberti, Alex S. Holehouse, Timothy T. Weil

## Abstract

Biomolecular condensates that form via liquid-liquid phase separation can exhibit diverse physical states. Despite considerable progress, the relevance of condensate physical states for *in vivo* biological function remains limited. Here, we investigated the physical properties of *in vivo* processing bodies (P bodies) and their impact on mRNA storage in mature *Drosophila* oocytes. We show that the conserved DEAD-box RNA helicase Me31B forms P body condensates which adopt a less dynamic, arrested physical state. We demonstrate that structurally distinct proteins and hydrophobic and electrostatic interactions, together with RNA and intrinsically disordered regions, regulate the physical properties of P bodies. Finally, using live imaging, we show that the arrested state of P bodies is required to prevent the premature release of *bicoid* (*bcd*) mRNA, a body axis determinant, and that P body dissolution leads to *bcd* release. Together, this work establishes a role for arrested states of biomolecular condensates in regulating cellular function in a developing organism.

## INTRODUCTION

Many biochemical reactions in the cytoplasm of eukaryotic cells require regulation in space and time. The organization of specific reactions in the dense cytoplasmic environment is achieved through membrane-bound and membrane-less organelles. Classic membrane-bound organelles such as the nucleus and endoplasmic reticulum are stable micro-environments enclosed by membranes. However, membrane-less organelles such as stress granules, processing bodies (P bodies) and nuclear bodies have been shown to also provide an essential, dynamic level of cellular organization (Banani *et al*., 2017; Shin and Brangwynne, 2017; Boeynaems *et al*., 2018). Typically composed of ribonucleic acids (RNA) and proteins, the assembly of many membrane-less organelles is well-described via a spontaneous demixing process referred to as liquid-liquid phase separation (LLPS) (Brangwynne *et al*., 2009; Li *et al*., 2012; Hubstenberger *et al*., 2013; Wang *et al*., 2014; Nott *et al*., 2015; Zhang *et al*., 2015; Feric *et al*., 2016). In its simplest instantiation, LLPS occurs when a specific set of one or more macromolecules condense into a distinct liquid-like phase that is relatively enriched for those macromolecules in comparison to the surrounding and coexisting phase (e.g., cytoplasm or nucleoplasm) (Hyman, Weber and Jülicher, 2014; Brangwynne, Tompa and Pappu, 2015; Banani *et al*., 2017; Lyon, Peeples and Rosen, 2021). More generally, the designation *biomolecular condensates* has emerged as an overarching term to describe cellular assemblies characterized by the non-stoichiometric concentration of biomacromolecules, of which membrane-less organelles are one such example.

Ribonucleoprotein (RNP) complexes are an abundant and conserved class of biomolecular condensates. Found both in the cytoplasm and the nucleus, RNP condensates exhibit a wide range of physical states ranging from dynamic liquids to stable solids (Weber and Brangwynne, 2012; Kroschwald *et al*., 2015, 2018; Weber, 2017; Woodruff *et al*., 2017). Some examples of diverse physical states observed *in vivo* include liquid-like P granules in *C. elegans* embryos (Brangwynne *et al*., 2009; Wang *et al*., 2014); viscous, semi-liquid nucleolus in *Xenopus laevis* oocytes (Brangwynne, Mitchison and Hyman, 2011; Feric *et al*., 2016; Mitrea *et al*., 2016); and solid-like Balbiani bodies in the cytoplasm of vertebrate oocytes (Boke *et al*., 2016). Despite considerable progress in our understanding of biomolecular condensates, the relationship between condensate physical state and *in vivo* function remains poorly understood. More broadly, while the study of metazoan condensates *in vitro* and in cell culture has been instrumental in our modern understanding of their role in cellular organization and function, *in vivo* approaches are much less common. RNP condensates are often linked with localized translation which enables cells to spatiotemporally regulate protein synthesis through localization, storage and translational control of stored mRNAs (Medioni, Mowry and Besse, 2012). Specialized cells, including neurons and eggs, often rely on this mode of post transcriptional regulation to control gene expression (Kloc and Etkin, 2005; Jung *et al*., 2014). Specifically, in transcriptionally inactive oocytes, such as *Drosophila melanogaster*, prolonged storage and translational control of maternally deposited transcripts are required for body axes patterning (Tadros and Lipshitz, 2009; Lasko, 2012).

One mechanism for regulating RNA storage and translational control involves P bodies, an evolutionarily conserved class of cytoplasmic biomolecular condensates (Andrei *et al*., 2005; Kedersha *et al*., 2005; Eulalio *et al*., 2007; Parker and Sheth, 2007; Buchan, Buchan and Buchan, 2014; Hubstenberger *et al*., 2017; Luo, Na and Slavoff, 2018). Previous work on P bodies has highlighted their roles in various aspects of RNA metabolism, including RNA storage and translational repression since P bodies are devoid of ribosomes (Weil *et al*., 2012; Hubstenberger *et al*., 2017). A conserved component of P bodies is the ATP-dependent DEAD box RNA helicase human DDX6, which is conserved across yeast (Dhh1), *C. elegans* (CGH-1), and *Drosophila* (Maternal expression at 31B (Me31B)). Me31B is required for early *Drosophila* development and estimated to be present at concentrations of ∼7.5 μM in the egg (Götze *et al*., 2017). During oogenesis, Me31B is known to associate with and differentially regulate several axis-patterning maternal mRNAs (Nakamura *et al*., 2001; Weil *et al*., 2012). The dorsoventral determinant *gurken* mRNA is translated early in oogenesis, while *bicoid* (*bcd*) mRNA is repressed and retained in the oocyte until egg activation, a conserved process that is required before entry into embryogenesis (Tadros and Lipshitz, 2009; Weil *et al*., 2012; Kaneuchi *et al*., 2015; York-andersen *et al*., 2015). Failure to regulate these, and many other mRNAs can lead to severe developmental defects (Lasko, 2012). However, the mechanisms that underlie how transcripts are maintained and translationally controlled by P bodies is not well understood.

To examine the *in vivo* basis of mRNA regulation, we use time-lapse fluorescence microscopy, genetics and pharmacological treatments to investigate the physical properties and *in vivo* function of P bodies in the mature *Drosophila* oocyte. We find that P body condensates adopt an arrested physical state in the mature oocyte. We show that P body integrity requires weak electrostatic and hydrophobic interactions, along with RNA and the actin cytoskeleton. Using *in silico* and *in vitro* approaches, we demonstrate that intrinsically disordered regions (IDRs) regulate Me31B condensation and their physical properties. We also find that the highly disordered protein Trailer hitch (Tral) is required for the assembly and organization of P bodies. Finally, we demonstrate that the condensed and arrested state of P bodies prevents the release of *bicoid (bcd)* mRNA, highlighting an important cellular function of P bodies in *Drosophila* and an *in vivo* role for P body physical state. Based on these results and other analogous work, we propose that less dynamic, arrested physical states are a universal feature of RNP condensates in specialized cells to temporally regulate RNAs in response to cellular and developmental cues.

## RESULTS

### Me31B forms less dynamic, arrested P body condensates in the mature oocyte

Maternal mRNAs are thought to be stored and regulated by P bodies throughout oogenesis (Nakamura *et al*., 2001; Lin *et al*., 2008; Weil *et al*., 2012). Egg activation of the mature oocyte, which is the final stage of oogenesis, results in the release of stored mRNAs and subsequent translation (Tadros and Lipshitz, 2009; Krauchunas and Wolfner, 2013). However, a mechanistic understanding of how the physical state of P bodies regulates mRNA remains unclear. To address this, we isolated living mature oocytes from female *Drosophila* (Figure 1A) and utilized the conserved DEAD-box RNA helicase, Me31B, to visualize P bodies (Figure 1B). Live imaging of Me31B (Me31B::GFP) revealed that P bodies are typically micron sized condensates with varying morphology and sizes (Figure 1C). We also find that most P bodies have internal subdomains indicating that they are heterogeneously organized (Figure 1D). Quantification of P body aspect ratios over time suggests that P bodies have predominantly irregular morphologies compared to liquid-like condensates which are typically spherical (Figure 1E). Additionally, P bodies undergo continuous morphological rearrangement, with most progressing from irregular to more rounded shapes (Figure 2A). Aspect ratio analysis of individual condensates followed over time showed rearrangement of P bodies relaxing from an amorphous to a round shape in 30 minutes (Figure 2A), more slowly than many previously studied liquid-like condensates (Brangwynne *et al*., 2009; Feric *et al*., 2016). Although P bodies undergo fusion and fission events (Figure 2B, 2B’, and S1), which are a hallmark of liquid-like condensates, the longer timescale of these events indicate that P bodies in the mature *Drosophila* oocyte are less dynamic compared to known liquid-like condensates such as P granules and stress granules (Brangwynne *et al*., 2009; Patel *et al*., 2015).

**Figure 1:**
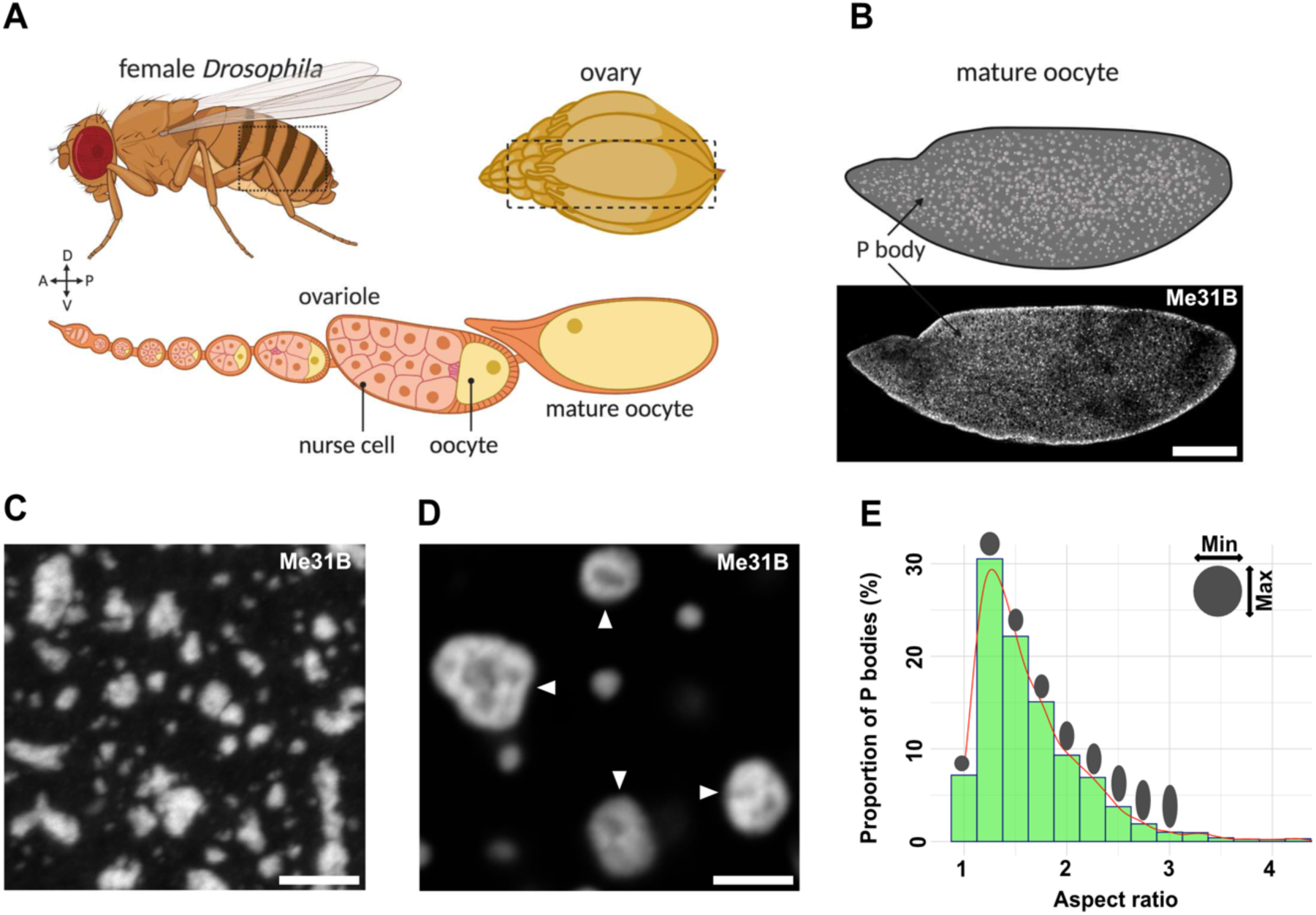
Me31B forms heterogeneous P body condensates in the mature oocyte. **(A)** Schematic of a *Drosophila* female, ovary and ovariole. Each female contains two ovaries that are each comprised of 16-18 ovarioles. Each ovariole can be thought of as an assembly line for the production of mature oocytes. The oocyte is supported by a collection of nurse cells until the late stages of oogenesis. Created with BioRender.com **(B-E)** Mature oocyte expressing Me31B::GFP. **(B)** Cartoon depicting P body distribution in the mature oocyte and confocal image of a whole mature oocyte showing P bodies throughout the cytoplasm. The concentration of P bodies at the cortex is in part due to this being a cross section image. **(C)** Increased magnification reveals P bodies exhibit diverse morphologies and sizes. Maximum projection 10 µm. **(D)** Representative image of P bodies exhibiting multiple subdomains (white arrowheads) indicative of heterogeneous internal organization. **(E)** Aspect ratio analysis of individual P bodies (>1 µm) showing an uneven range of P body morphology. Scale bar = 100 µm (B), 5 µm (C,D).

**Figure 2:**
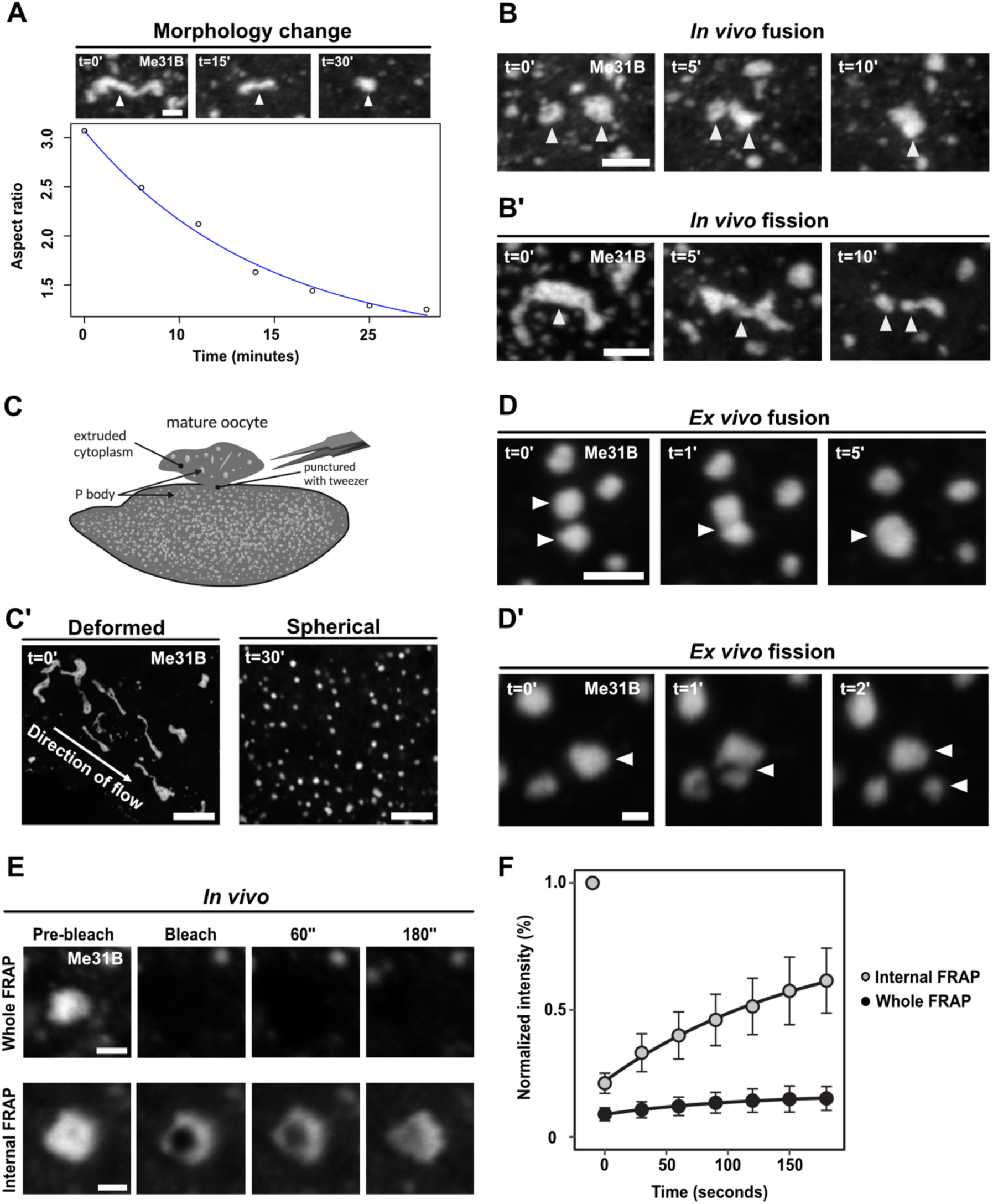
P bodies adopt a less dynamic and arrested physical state. **(A, B, B’, C’, D, D’, E)** Mature oocyte expressing Me31B::GFP. **(A)** Time series of a P body displaying elastic behaviour, starting in an extended state (t=0’) and subsequently relaxing towards a spherical morphology (t=30’). Plot of individual P body (n = 10) A.R over time showing relaxation from extended (A.R∼3) to spherical morphology (A.R∼1). **(B)** Time series of two *in vivo* P bodies undergoing coalescence (white arrowheads). (**B’**) Time series of a single *in vivo* P body undergoing fission to form two distinct condensates (white arrowheads). **(C)** Cartoon depicting cytoplasmic extrusion of P bodies into halocarbon oil (*ex vivo*) induced by puncturing the outer membrane of the mature oocyte. Created with BioRender.com **(C’)** Confocal image of *ex vivo* P bodies displaying stretched elastic morphologies shortly after extrusion (t=0’). Over time, extruded P bodies relax into homogeneous spherical condensates (t=30’). **(D)** Time series of *ex vivo* P bodies undergoing coalescence (white arrowheads). **(D’)** Time series of *ex vivo* extruded P bodies undergoing fission (white arrowheads). **(E)** Time series of whole FRAP of P body shows minimal recovery, while internal FRAP of P body shows increased recovery of Me31B fluorescence. **(F)** P body recovery profiles after whole FRAP and internal FRAP (n = 20 P bodies for whole FRAP and n = 24 P bodies for internal FRAP). Scale bar = 2.5 µm (A), 5 µm (B,B’,D,D’), 10 µm (C’), 1.5 µm (E).

The cytoplasm of the mature oocyte is packed with yolk granules, organelles, and complex cytoskeletal structures. To determine if the slow P body dynamics are an intrinsic property or dependent on the oocyte cytoplasmic environment, we developed an *ex vivo* assay whereby we extrude the cytoplasm into halocarbon oil (Figure 2C). Importantly, this approach does not promote P body dissolution, rather, extruded P bodies initially exhibit irregular morphologies but become spherical with time (Figure 2C’). Consistent with *in vivo* observations, we also show extruded P bodies undergo fusion and fission events over longer timescales (Figure 2D and 2D’). Taken together our data suggests that P bodies are slowly rearranging viscoelastic condensates, and that this physical state is inherent to P bodies from mature *Drosophila* oocytes.

We next performed Fluorescence Recovery After Photobleaching (FRAP) on whole P bodies (whole FRAP) to examine the mobility of Me31B between the cytoplasm and P body. This analysis revealed that Me31B localized to P bodies exhibited limited or no recovery (Figure 2E and 2F). Due to their inability to exchange Me31B with the cytoplasm, we refer to this as the arrested state of P bodies. To further explore Me31B dynamics, we tested if Me31B can rearrange within P bodies by observing the recovery of Me31B fluorescence after photobleaching within a region inside the P body (internal FRAP) (Figure 2E and 2F). Measurements revealed considerable recovery that progresses from the periphery to the center of the P body (Figure S2A). Despite exhibiting a higher mobile fraction than whole FRAP, the rate of recovery (Figure S2B) indicates that Me31B exchanges slower internally compared to liquid-like condensates. Overall, these data show that P bodies in the mature oocyte adopt a less dynamic and arrested physical state.

### Weak intermolecular, cytoskeletal and RNA interactions regulate P physical properties

Previous work has shown that activation of the mature oocyte results in an influx of monovalent and divalent ions, release of stored mRNAs, and reorganization of the actin cytoskeleton (Kaneuchi *et al*., 2015; York-andersen *et al*., 2015; Andersen *et al*., 2020). We therefore wondered if these factors could regulate the physical properties of P bodies in the mature oocyte, prior to egg activation.

Various molecular interactions have been shown to contribute to RNP condensation including electrostatic, cation-pi and hydrophobic interactions (Kato *et al*., 2012; Brangwynne, Tompa and Pappu, 2015; Nott *et al*., 2015; Pak *et al*., 2016; Riback *et al*., 2017; Murthy *et al*., 2019; Dzuricky *et al*., 2020). The interactions that are thought to drive the assembly of P bodies can be interpreted through the lens of a pseudo-two component phase diagram (Figure 3A). In particular, by changing the solution conditions to weaken the interactions that contribute to P body assembly, theory and simulations predict an increase in internal mobility and more spherical condensates, as shown previously for protein-RNA condensates (Boeynaems *et al*., 2019).

**Figure 3:**
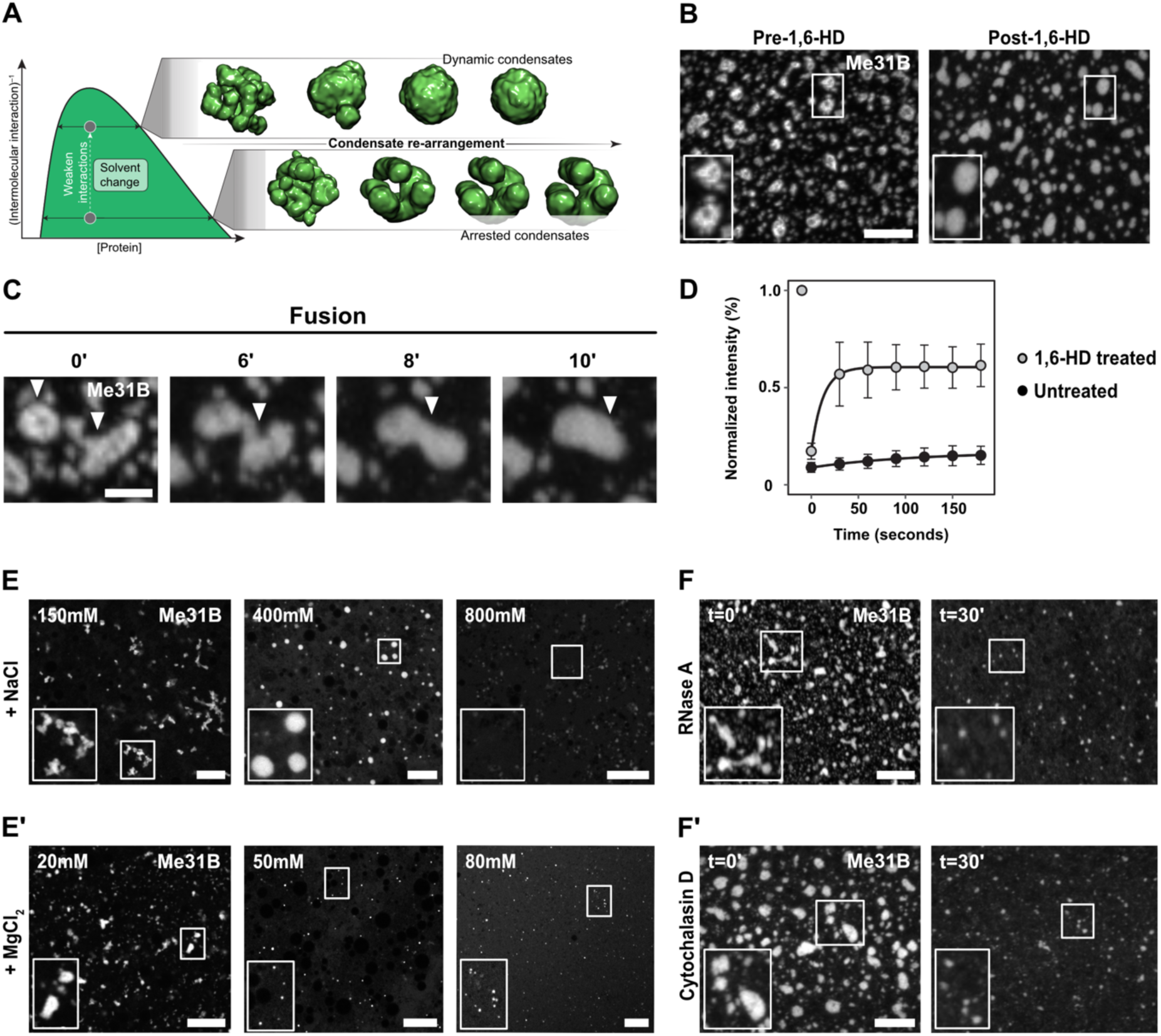
P body physical properties are regulated by hydrophobic and electrostatic interactions along with RNA and actin. **(A)** Schematized phase diagram in which protein concentration extends along the x-axis whereas molecular interaction strength extends across the y-axis. Inset shows snapshots from coarse-grained simulations performed at distinct position along the y-axis. Condensate morphology is dependent on intramolecular interaction strength, such that weak intermolecular interactions lead to spherical condensates, while strong intermolecular interactions lead to kinetically arrested amorphous condensates. **(B,C,E-F’)** Mature oocyte expressing Me31B::GFP. **(B)** Addition of 5% 1,6-HD causes P bodies to transform from amorphous to spherical morphology. Furthermore, addition of 1,6-HD results in the loss of internal heterogeneity of P bodies. Maximum projection 5 µm. **(C)** Time series shows two P bodies undergoing coalescence following the addition of 1,6-HD. Maximum projection 5 µm. **(D)** Whole FRAP recovery profile of 1,6-HD treated P bodies (n = 12) showing rapid fluorescence recovery compared to untreated P bodies (n = 24). **(E)** Addition of varying concentrations of NaCl results in diverse physical states of extruded P bodies ranging from sticky (150 mM) to liquid-like (400 mM) and diffuse state (800 mM). **(E’)** Treatment with MgCl_2_ results in the dissociation of extruded P bodies at concentrations significantly lower than NaCl. **(F)** Treatment with 500 ng/µl RNase A (degrades RNA) or **(F’)** 10 µg/µl cytochalasin-D (depolymerizes actin) causes P body dissociation, resulting in smaller condensates. Maximum projection 10 µm. Scale bar = 5 µm (B,C,E,E’,F,F’)

To test if hydrophobic interactions are required for *in vivo* P body integrity, we treated mature oocytes with the aliphatic alcohol 1,6-hexanediol (1,6-HD), a compound identified originally in the context of attenuating hydrophobic interactions (Ribbeck and Go, 2002; Patel *et al*., 2007). Addition of 1,6-HD resulted in the transformation of P bodies from irregular to spherical morphologies and an increase in condensate fusion events (Figure 3B, 3C, and S3A). These results support a model in which 1,6-HD weakens the intermolecular interactions that contribute to P body integrity. To further test if the disruption of hydrophobic interactions leads to a transition from an arrested to a more dynamic state, we performed whole FRAP on 1,6-HD treated P bodies. Consistent with our model, 1,6-HD treated P bodies exhibit rapid and sustained recovery (Figure 3D, S3B, and S3C). Taken together, our results suggest that hydrophobic interactions contribute to regulating the arrested state of P bodies.

We next examined if electrostatic interactions contribute to P body physical properties by testing the impact of monovalent (NaCl) or divalent salt (MgCl_2_). At low concentrations of NaCl, P bodies assemble into clusters while at high concentrations they dissociate (Figure 3E). However, at a specific range of concentrations (300 – 600 mM), P bodies adopt spherical morphologies, consistent with a more dynamic state. These results are supportive of a model in which electrostatic interactions, like hydrophobic interactions, play a role in dictating physical properties and can be tuned up or down by decreasing or increasing the monovalent salt concentrations, respectively.

Interestingly, the addition of 20 mM MgCl_2_ had no apparent effect on P body integrity, yet a small increase to concentrations as low as 50 mM MgCl_2_ results in their complete dissociation (Figure 3E’). This relative sensitivity to divalent cations implies an effect beyond simply ionic strength. Collectively these data suggest that changes in salt concentration can alter P body integrity, likely via both ion-specific effects and electrostatic screening. This is consistent with the morphology and state of P bodies that we observe following *ex vivo* egg activation or in the early embryo (York-andersen *et al*., 2015).

Given the importance of electrostatic interactions, we asked if P body integrity was regulated exclusively by protein-protein interactions, or if protein-RNA interactions also contribute. To test this, we treated mature oocytes with RNase A, which leads to P body dissociation over a time course of 30 minutes (Figure 3F). This result reveals that that *in vivo* P body integrity depends, at least in part, on protein:RNA interactions but this does exclude a contribution from protein-protein interactions.

Finally, to examine the role of actin in regulating P body integrity, mature oocytes were treated with cytochalasin D, a commonly used actin depolymerizing agent. This treatment resulted in the dissociation of P bodies in 30 minutes, consistent with our data from *ex vivo* egg activation (Figure 3F’). Taken together, these results indicate that multiple factors regulate P body integrity and morphology in the mature oocyte.

### IDRs regulate the physical state of Me31B condensates *in vitro*

Having established a role for multiple external factors in the regulation of P body integrity, we next asked how the Me31B protein may be contributing to P body physical state. Me31B contains an ATP-binding domain and folded helicase domain, flanked by short N- and C-terminal IDRs (Figure 4A). While the function of the helicase domain is well known, the function of the disordered regions is rather unclear. Since Me31B is an essential *in vivo* protein, we adopted an *in vitro* approach to examine the role of these disordered regions.

**Figure 4:**
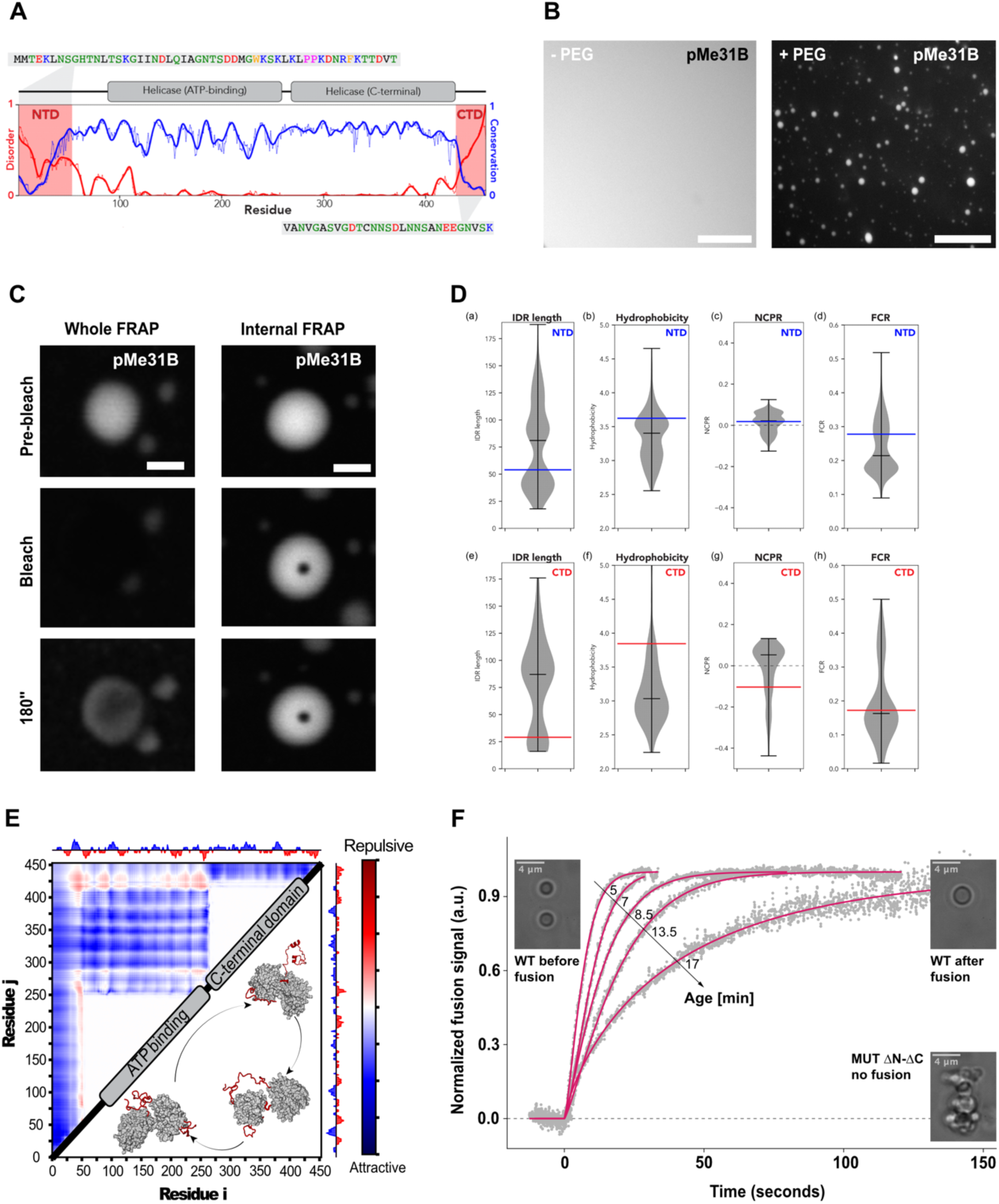
Deletion of IDRs results in the aggregation of Me31B condensates *in vitro*. **(A)** Purified GFP-Me31B (pMe31B) at 7.5 µM is diffuse on its own, but forms phase separated spherical condensates in the presence of 1% PEG. Maximum projection 5µm. **(B)** Time series of pMe31B condensates subjected to FRAP experiments. Whole P body photobleaching shows moderate fluorescence recovery, while internal FRAP shows no recovery. **(C)** Violin plots quantify density of IDR length (a/e), hydrophobicity (b/f), Net charge per residue (c/g) and fraction of charged residues (d/h) for the N-terminal IDRs (a-d) or C-terminal IDRs (e-h). Blue or red bars define the associated value for the Me31B IDR in the N- or C-terminal IDR, respectively. **(D)** Overview of disordered, conservation, and domain architecture for Me31B. Conservation calculated across 566 orthologous sequences. CTD and NTD sequences are highlighted, with an atomistic model of the full-length protein shown below. **(E)** Summary of all-atom simulations. Normalized inter-residue distance is shown, with cooler colors reflecting attractive interactions and warmer colors reflecting repulsive interactions. Normalized distances are calculated based on the expected distance for a self-avoid polymer model. Both the NTD and CTD engage directly with the folded domains in a distributed and transient manner. Interactions are relatively uniform across the folded domain surface. **(F)** Fusion of pMe31B condensates (magenta) at different time points post condensation, quantified by dual-trap optical tweezers. pMe31B ΔN-ΔC condensates (dashed line) do not fuse and rapidly aggregate with each other. Scale bar = 5µm (A,F), 2.5µm (B).

First, we tested if the purified recombinant Me31B (GFP-Me31B) can undergo phase separation *in vitro*. While Me31B is diffuse at physiological protein concentrations (7.5 μM), upon the addition of a small amount of crowding agent (1% PEG) Me31B undergoes phase separation to form condensates (Figure 4B). We reproduced Me31B condensation using a secondary crowding agent (1% Ficoll), demonstrating that Me31B phase separation does not result from the specific chemical properties of the used crowding agent (Figure S4A). Time lapse imaging revealed that Me31B initially forms condensates with highly dynamic properties, as evidenced by an increase in condensate size (Figure S4B). However, over time these condensates become less dynamic, as the apparent fusion kinetics of Me31B condensates slows as a function of time (Figure S4C). To test whether or not the Me31B molecules in the initial condensates are mobile, we performed both whole FRAP and internal FRAP on freshly formed condensates. To our surprise, these showed little or no recovery after photobleaching, indicating that Me31B condensates are present in an arrested physical state similar to *in vivo* P bodies (Figure 4C).

It has been suggested that the assembly and properties of RNP condensates is regulated by IDRs (Lin, David S.W. Protter, *et al*., 2015; Martin and Mittag, 2018; Martin *et al*., 2020). Previous work has shown that the sequence length and position of IDRs varied greatly between DDX6 and Dhh1 (Hondele *et al*., 2019). To examine the extent of variation across the DDX6 family of proteins, we performed a sequence analysis of Me31B orthologs. This revealed that folded domains are highly conserved while IDR length and sequence varied substantially (Figure 4D).

To better understand how the IDRs might contribute to function, we performed all-atom simulations of full-length Me31B. Simulations revealed that both IDRs adopt a heterogeneous ensemble of states (Figure 4E). Interestingly, both N- and C-terminal IDRs interacted transiently and relatively non-specifically with the surface of the folded domains. These contacts were mediated through electrostatic and hydrophobic interactions (Figure 4F, S4D, and S4E). Rather than acting as drivers of self-assembly, our simulations suggest the possibility that IDRs play a modulatory role.

To test for the modulatory influence of IDRs in Me31B phase separation, we next purified recombinant Me31B with both IDRs deleted (Me31BΔN-ΔC). Consistent with the interpretation from our simulations, Me31BΔN-ΔC rapidly self-assembled into solid-like aggregates (Figure 4F and S4F). These results demonstrate that protein-protein interactions can be established independently of the IDRs and suggests that the folded domain plays an important role in driving self-assembly. Importantly, our results demonstrate that the IDRs tune the physical properties of Me31B condensates, attenuating the strong interactions established among the interacting folded domains.

Taken together, our results show that recombinant Me31B is prone to form condensates with a less dynamic and arrested physical state, consistent with *in vivo* data. Importantly, the short IDRs from Me31B function to modulate the condensate physical state, suggesting that the interactions that drive Me31B self-assembly originate from the folded domains.

### Tral is key to regulating organization of P bodies in the mature oocyte

In addition to Me31B, several other proteins localize to, or are found to be enriched within P bodies (Lin *et al*., 2008). Given the importance of disordered regions within Me31B, we hypothesized that intrinsically disordered proteins (IDPs) within P bodies could potentially act as lubricants to regulate P body assembly and organization through interactions with structured proteins. To test this, we first performed disorder prediction across the set of known P body proteins to estimate the proportion of structured versus disordered regions (Figure 5A). Approximately 50% of proteins known to localize to P bodies in *Drosophila* contain long IDRs, highlighting the structural heterogeneity of components within P bodies. Among the proteins enriched with intrinsic disorder is Tral, a member of the LSM protein family (RAP55 in vertebrates, CAR-1 in *C. elegans*). Additionally, Tral is also known to interact directly with Me31B, function in *Drosophila* axis patterning (Bouveret, 2000; Monzo *et al*., 2006; Tritschler *et al*., 2007, 2008, 2009; Götze *et al*., 2017; Wang *et al*., 2017; Hara *et al*., 2018; McCambridge *et al*., 2020) and is predicted to be largely disordered with the exception of an N-terminal LSM domain (Figure 5B).

**Figure 5:**
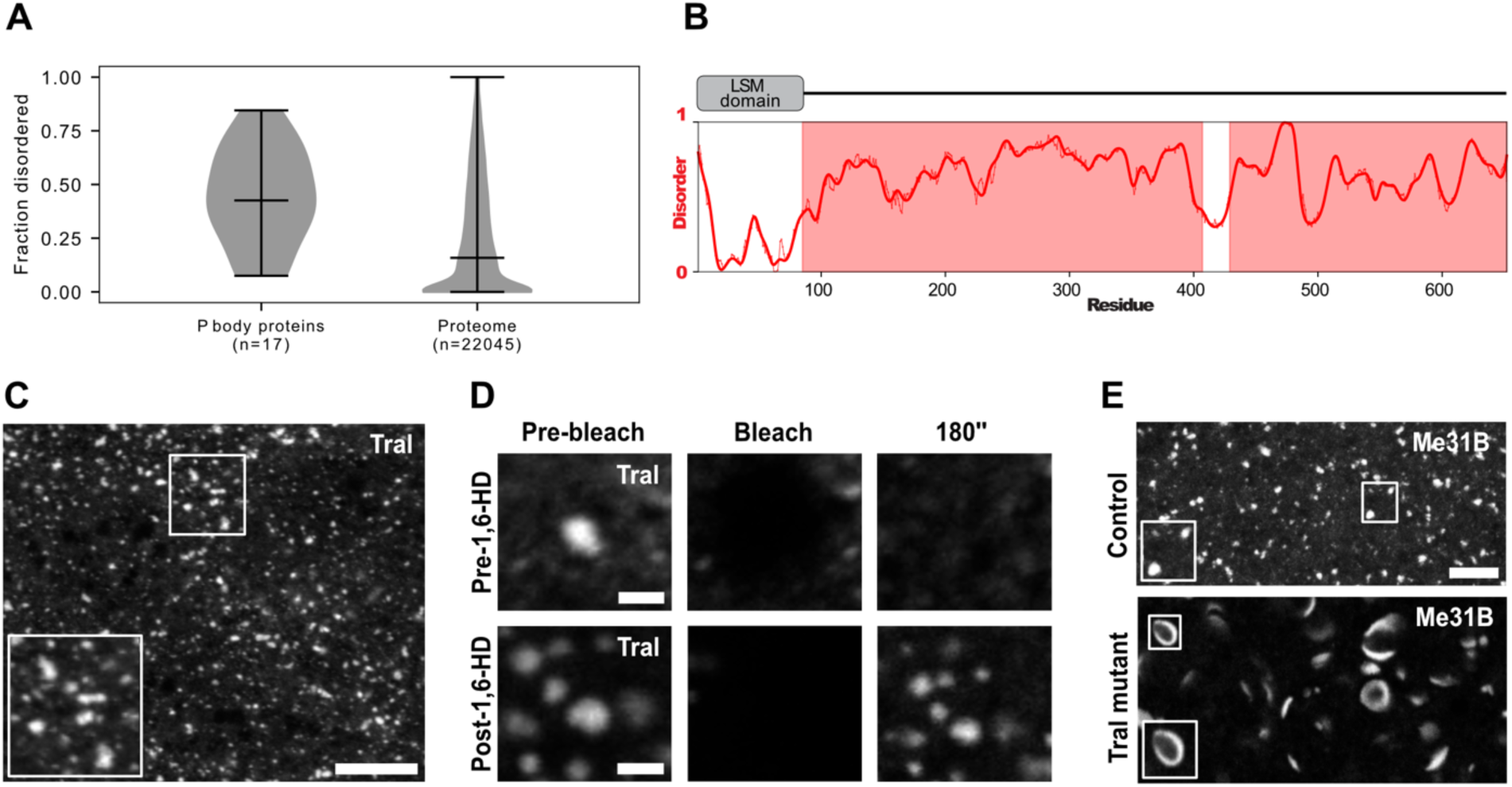
Absence of Tral alters P body morphology in the mature oocyte. **(A)** Comparison of fraction disorder in known *Drosophila* P body proteins (left) compared to whole *Drosophila* proteome (right). **(B)** Schematic of Tral domain architecture containing a structured LSM domain followed by a long stretch of highly disordered regions. **(C,D)** Mature oocyte expressing GFP::Tral. **(C)** Tral localizes to P body condensates with diverse morphologies and sizes, distributed throughout the oocyte cytoplasm. Maximum projection 7 µm. **(D)** Time series of FRAP experiments on GFP::Tral condensates before and after treatment with 1,6-HD. Prior to the addition of 1,6-HD, Tral condensates do not recover after photobleaching, however, Tral condensates treated with 1,6-HD show increased fluorescence recovery post photobleaching. **(E)** Representative mature oocytes expressing Me31B::GFP, control (*tral^1^* / *+* or *Df(3L)ED4483 / +*) displaying close to spherical P body condensates. In the absence of Tral (*tral^1^ / Df(3L)ED4483*), Me31B forms aberrant rod and doughnut shaped P body condensates. Maximum projection 5 µm. Scale bar = 5µm (C,E), 1.5µm (D).

We therefore tested the role of Tral in the regulation of P body assembly *in vivo*. In mature oocytes expressing GFP::Tral, like Me31B, Tral associates with P body condensates (Figure 5C). Despite being localized to the same condensate, different proteins do not need to follow equivalent dynamics (Boeynaems *et al*., 2019). We therefore asked whether TraI shows similar *in vivo* behaviors to Me31B, which would suggest that these two proteins are associated with the same physical state. Indeed, despite being structurally distinct from Me31B, whole FRAP and 1,6-HD experiments on Tral were consistent with our results for Me31B (Figure 5D). This supports a model in which Me31B and TraI are strongly coupled within P bodies, likely through direct interaction or interaction with a common intermediate (e.g., RNA).

Work in arrested *C. elegans* oocytes has shown that the organization and properties of germline P bodies is regulated by orthologs of Tral (CAR-1) and Me31B (CHG-1), where loss of CGH-1 resulted in the formation of morphologically deformed P bodies with altered physical properties (Hubstenberger *et al*., 2013). Since Me31B is essential for *Drosophila* oogenesis, we tested if Tral is required to regulate Me31B labeled P bodies in the mature oocyte. Remarkably, in Tral mutants, P bodies have dramatically different morphologies and form rod and planar donut-shaped assemblies (Figure 5E), implying a gain of anisotropy in the underlying molecular arrangement of the condensate. The formation of apparently ordered (or partially ordered) assemblies is reminiscent of liquid-crystalline formation as observed in the synaptonemal complex or in specific mutants of the plant protein FLOE1 (Rog, Köhler and Dernburg, 2017; Dorone *et al*., 2020). These results suggest that despite being structurally distinct, Tral and Me31B contribute to the assembly and organization of P bodies likely through synergistic interactions.

### The condensed and arrested state of P bodies regulate *bcd* mRNA

Our data shows that P bodies in the mature *Drosophila* oocyte are present in a less dynamic, arrested physical state. This is in contrast to previous data in cells where related RNP condensates are shown to exhibit dynamic behavior adopting liquid-like states (Kroschwald *et al*., 2015). Since P bodies in the mature oocyte contain maternal mRNAs that are stored and translationally regulated over long periods, we hypothesized that the less dynamic and arrested physical state could be required for this function.

The anterior determinant *bcd* mRNA is a well-established example of long-term storage and localizes to P bodies in the mature oocyte (Figure 6A). To address if the condensed state of P bodies is required for the association with *bcd* mRNA, we first tested the change in P body and *bcd* mRNA distribution following egg activation. Using a previously established *ex vivo* egg activation assay, we show that both P bodies and *bcd* mRNA undergo rapid dispersion (Figure 6B). This result indicates that the transition of P bodies from a condensed to diffused state leads to the release of *bcd* mRNA. This finding is consistent with data arguing that translation of *bcd* mRNA only occurs after egg activation and when the mRNA is no longer inside of P bodies (Weil *et al*., 2012; Eichhorn *et al*., 2016).

**Figure 6:**
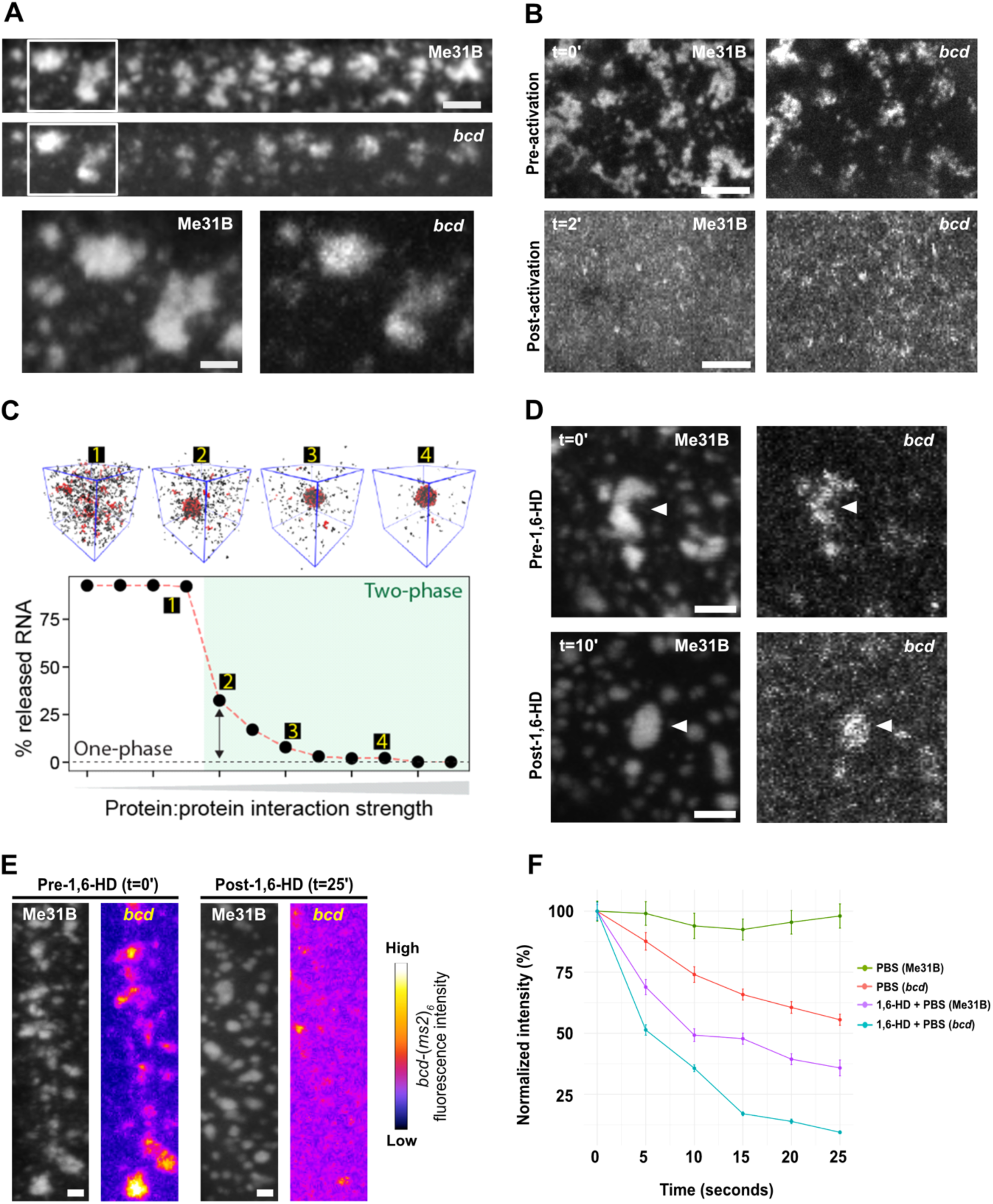
Altering P body physical state leads to premature loss of *bcd* association with P bodies. **(A,B,D,E)** Mature oocyte expressing Me31B::GFP, *hsp83-MCP-RFP* and *bcd-*(*ms2*)*_6_*. **(A)** High resolution image of *bcd* mRNA co-localizing with P bodies at the anterior region of the mature oocyte. Inset shows a zoomed in version of *bcd* mRNA and P body association. Maximum projection 5 µm. **(B)** Upon addition of activation buffer, P bodies and *bcd* mRNA simultaneously undergo dispersion from condensed (t=0**’**) to diffused state (t=2’). Maximum projection 5 µm. **(C)** Coarse-grained simulations performed with 50 RNA molecules 800 protein molecules in which condensate assembly is driven by both protein:protein and protein:RNA interactions. When condensates have formed, the fraction of free (dilute phase) RNA is determined by both protein:protein and protein:RNA interaction strength. **(D)** Addition of 1,6-HD causes P bodies and *bcd* mRNA to initially adopt a spherical morphology (t=10’), representative of a more dynamic physical state. Maximum projection 5 µm. **(E)** Extended exposure to 1,6-HD results in dispersion of *bcd* mRNAs whereas P bodies remain condensed (t=25’). Maximum projection 5 µm. **(F)** Quantification of P body and *bcd* mRNA fluorescence in the presence of PBS (n = 37 Me31B and *bcd* particles) or 1,6-HD (n = 71 Me31B and *bcd* particles). Scale bar = 2.5µm (A), 5µm (B,D), 2µm (E).

To further explore this phenomenon, we developed a simple coarse-grained model in which protein and RNA will co-assemble to form condensates *in silico* (Figure 6C and S5). In our model, protein and RNA molecules possess attractive protein:protein and protein:RNA interactions that form multi-component condensates through phase separation. Condensate stability depends on both the strength of protein-protein and protein-RNA interactions, such that over the concentration range examined both species are necessary for phase separation. In simulations where the protein-protein interaction strength is systematically weakened we observe a concomitant release of RNA into the dilute phase and loss of condensate integrity. These simulations predict that condensate integrity can be viewed as a proxy for RNA sequestration and storage. We therefore sought to test this prediction *in vivo*.

Since we can experimentally alter the physical state of P bodies with small molecules, we asked if an arrested state is necessary for the association of *bcd* mRNA with P bodies. Upon 1,6-HD treatment of mature oocytes, both P bodies and *bcd* mRNA initially adopt spherical morphologies, consistent with a loss of P body integrity and transition to a more dynamic state (Figure 6D). However, in line with our predictions, while P bodies remain condensed and spherical, *bcd* mRNA becomes diffuse and is no longer observed in P bodies (Figure 6E and 6F). Together, these data suggest that changing the state of P bodies, either experimentally or with a developmental cue, causes release and potentially translation of stored mRNAs.

## DISCUSSION

Over the last decade, biomolecular condensates have emerged as a key principle in cellular organization. While changes in condensate physical properties have been examined extensively *in vitro*, the *in vivo* importance and function of physical states has been much less explored. In this study, we focused on the physical properties that are likely to enable P bodies to regulate mRNA in mature *Drosophila* oocytes. We demonstrate that a combination of intrinsic (intermolecular interactions, presence of IDRs) and extrinsic (RNA, actin, disordered proteins) factors can regulate the integrity and the arrested physical state of P bodies, both of which allow regulation of *bcd* mRNA *in vivo*. Overall, we propose that condensates with less dynamic, arrested physical states offer a controlled mechanism for long-term storage of translationally repressed mRNAs.

While dynamic, liquid-like states have been observed for many biomolecular condensates, there is a growing repertoire of functionally important and dynamically arrested condensates (Brangwynne, Mitchison and Hyman, 2011; Hubstenberger *et al*., 2013; Boke *et al*., 2016; Woodruff *et al*., 2017). Balbiani bodies, for instance, adopt a solid-like physical state which is thought to facilitate prolonged storage of organelles and macromolecules in dormant vertebrate oocytes (Boke *et al*., 2016). Similarly, the nucleolus in *Xenopus laevis* oocytes is shown to exist in a highly viscous physical state under physiological conditions (Brangwynne, Mitchison and Hyman, 2011). P bodies in *Drosophila* eggs also exhibit a similar physical state which likely enables long term storage of maternal mRNAs by strongly inhibiting access to ribosomes in the cytoplasm. Analogous properties have been observed in the germline P bodies of arrested *C. elegans* oocytes (Hubstenberger *et al*., 2013), suggesting that less dynamic and arrested properties of RNP condensates could be an evolutionarily conserved mechanism to temporally regulate mRNAs essential for normal development. Importantly, dynamically arrested physical states of RNP condensates are likely not limited to egg cells but may be preserved across other specialized cell types such as neurons. For example, mRNAs stored and translationally repressed in neuronal RNP condensates are temporally translated in an activity induced manner at specific synapses, thereby influencing short-term or long-term memory (Rajasethupathy *et al*., 2009; Puthanveettil, 2013). While it is not clear how this translation is regulated, models suggest that RNP granules switch between dynamic liquid-like and solid-like states to facilitate differential translation control (Majumdar *et al*., 2012; Sudhakaran and Ramaswami, 2017; Bakthavachalu *et al*., 2018). Beyond RNPs, in *A. thaliana,* dynamically arrested cytoplasmic condensates rewire transcription by sequestering a subset of the auxin-responsive transcription factor, preventing its nuclear function (Powers *et al*., 2019).

One striking observation is the influence of disordered regions and/or proteins in regulating the physical state of P body condensates. Conventional wisdom posits that IDRs contribute weak multivalent interactions that are essential for phase separation. In support of this model, many proteins that undergo phase separation contain IDRs that are necessary and sufficient for assembly (Lin, David S.W. Protter, *et al*., 2015; Protter *et al*., 2018). Our results offer an alternative model; rather than driving assembly, IDRs may also function to modulate and tune interactions between folded domains, potentially providing a local lubricant that counteracts strong multivalent interactions driven by adhesive interaction sites on folded domains. In the absence of IDRs, the folded domains from Me31B undergo irreversible aggregation. As such, the IDRs provide evolutionarily malleable modules to tune the physical state of condensates. This model echoes prior work on the yeast prion protein Sup35, where the loss of N-terminal disordered regions lead to robust aggregation of the folded C-terminal domain, while the full-length protein rapidly assembles into dynamic condensates that rapidly mature into gel-like assemblies (Franzmann *et al*., 2018). As such, we speculate that the disordered regions across Me31B orthologs (and indeed more generally DDX helicases) may have emerged to modulate the physical state of the resulting RNP complexes that form, be they micron-scale membrane-less organelles such as P bodies, or mesoscopic clusters of protein and RNA.

One other key determinant that regulates biomolecular condensates is multivalency. Condensates such as P bodies contain hundreds of diverse RNP components which serve as a major source of multivalent interactions that control their integrity and physical properties. While structured and disordered RNA-binding proteins have been investigated previously whether or how they influence the overall property of condensates *in vivo*, was unclear. Using Me31B and Tral, our data indicates that structurally distinct proteins synergistically interact to regulate the assembly and organization of P bodies during *Drosophila* oogenesis. Despite being structurally different to Me31B, Tral exhibits strikingly similar physical properties as Me31B when examined in P body condensates. However, loss of Tral results in structurally deformed and physically altered P bodies. These data also agree with observations reported for Tral and Me31B orthologues in arrested *C. elegans* oocytes (Hubstenberger *et al*., 2013) suggesting that despite sequence and structural divergence, the underlying molecular and physical interactions between components of RNP condensates may be evolutionarily conserved.

Prior to this work, it was unclear how stored mRNAs could be subjected to differential release and translation at distinct developmental stages without obviously disrupting the integrity of P bodies. As demonstrated for several RNA binding proteins, the assembly and physical properties of RNP condensates are largely regulated by weak intermolecular forces. We show that the physical properties of P bodies in the mature oocyte are disrupted by both 1,6-HD and salt, implicating a role for hydrophobic and electrostatic interactions in P body integrity. We speculate that modulating the strengths of different interactions allows P bodies to adjust their physical state, thereby facilitating fine-tuned control of mRNA storage and subsequent translation in response to developmental cues.

In summary, we describe the less dynamic, arrested physical state of Me31B labeled P body condensates and their potential role in regulating the storage of *bcd* mRNA in the mature *Drosophila* oocyte. The ability of weak intermolecular interactions, modular protein regions and cellular factors to regulate condensate physical state provides an elegant mechanism for P bodies to respond to cellular and developmental cues (Figure 7). We predict that such physical states are likely a universal feature of RNP condensates in specialized cells that require long term translational regulation of stored mRNAs.

**Figure 7:**
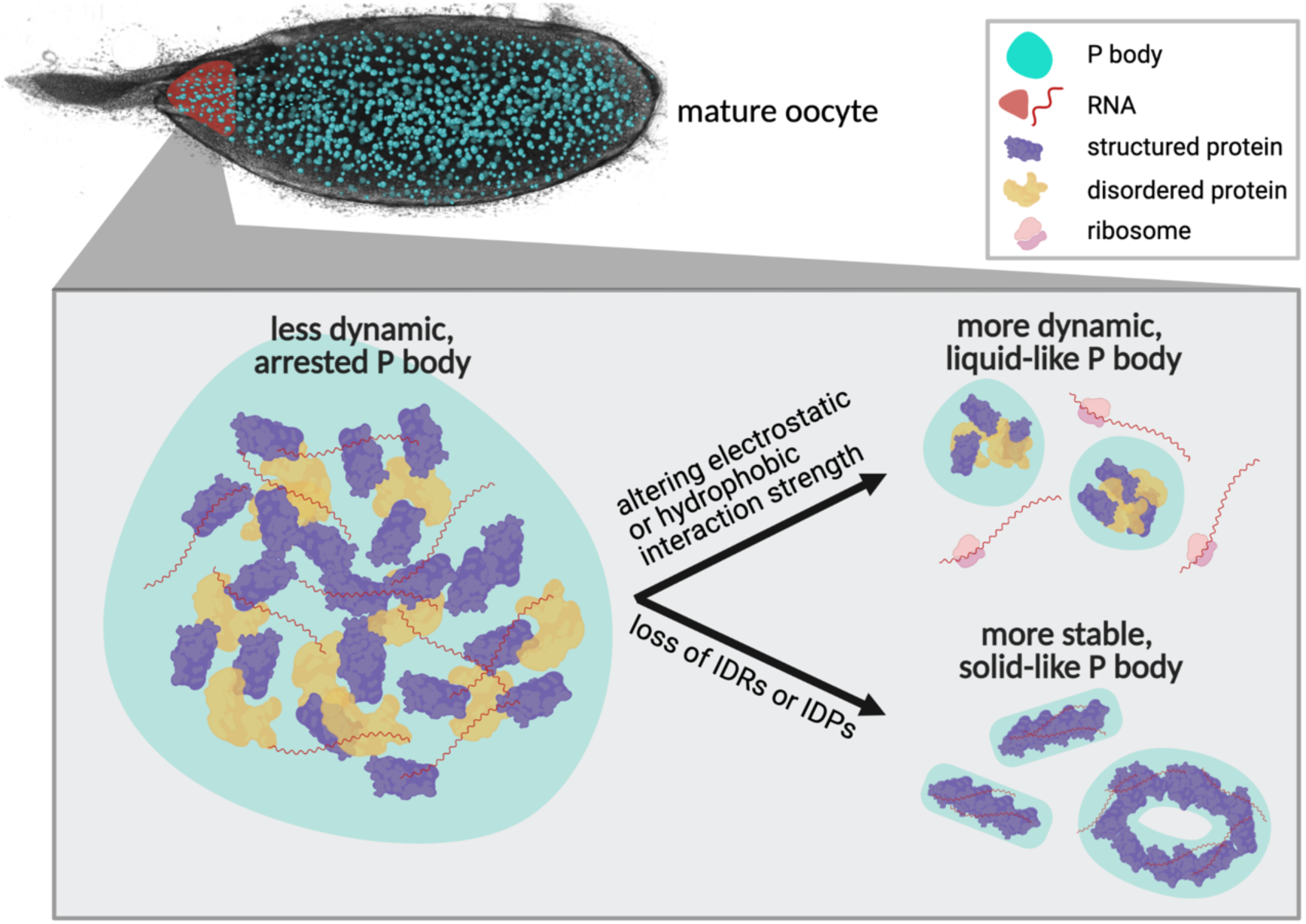
Regulation of P bodies in the mature *Drosophila* oocyte. P bodies (cyan) distributed throughout the mature *Drosophila* oocyte adopt a less dynamic, arrested physical state. The assembly, organization and physical properties of P bodies are regulated by synergistic interactions between structured (purple) and disordered proteins (yellow), as well as RNAs, predominantly via weak intermolecular hydrophobic and electrostatic interactions. Altering the strength of these interactions can lead to more dynamic, liquid-like P bodies and result in the release, and subsequent translation, of stored mRNAs (red). Alternatively, loss of IDRs or IDPs results in the formation of morphologically aberrant, solid-like protein-RNA aggregates which likely interfere with mRNA storage and translational regulation. Created with BioRender.com

---

**Figure S1:**
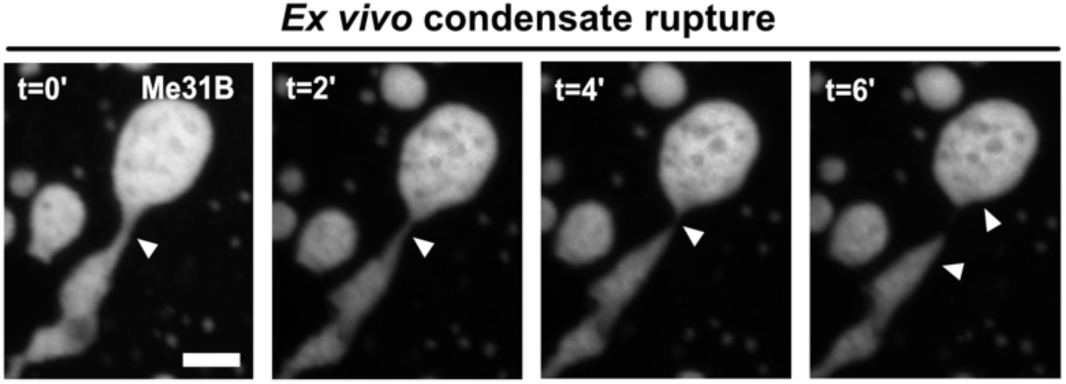
Extruded P bodies display rupture behavior. Time series of P bodies extruded from the mature oocyte are first held together by a transient ‘bridge’ after fusion. The P bodies then undergo a process of ‘pinching off’ of the bridge (white arrowheads point to the region of rupture). The two P bodies resorb within a minute of bridge rupturing. Scale bar: 5µm.

**Figure S2:**
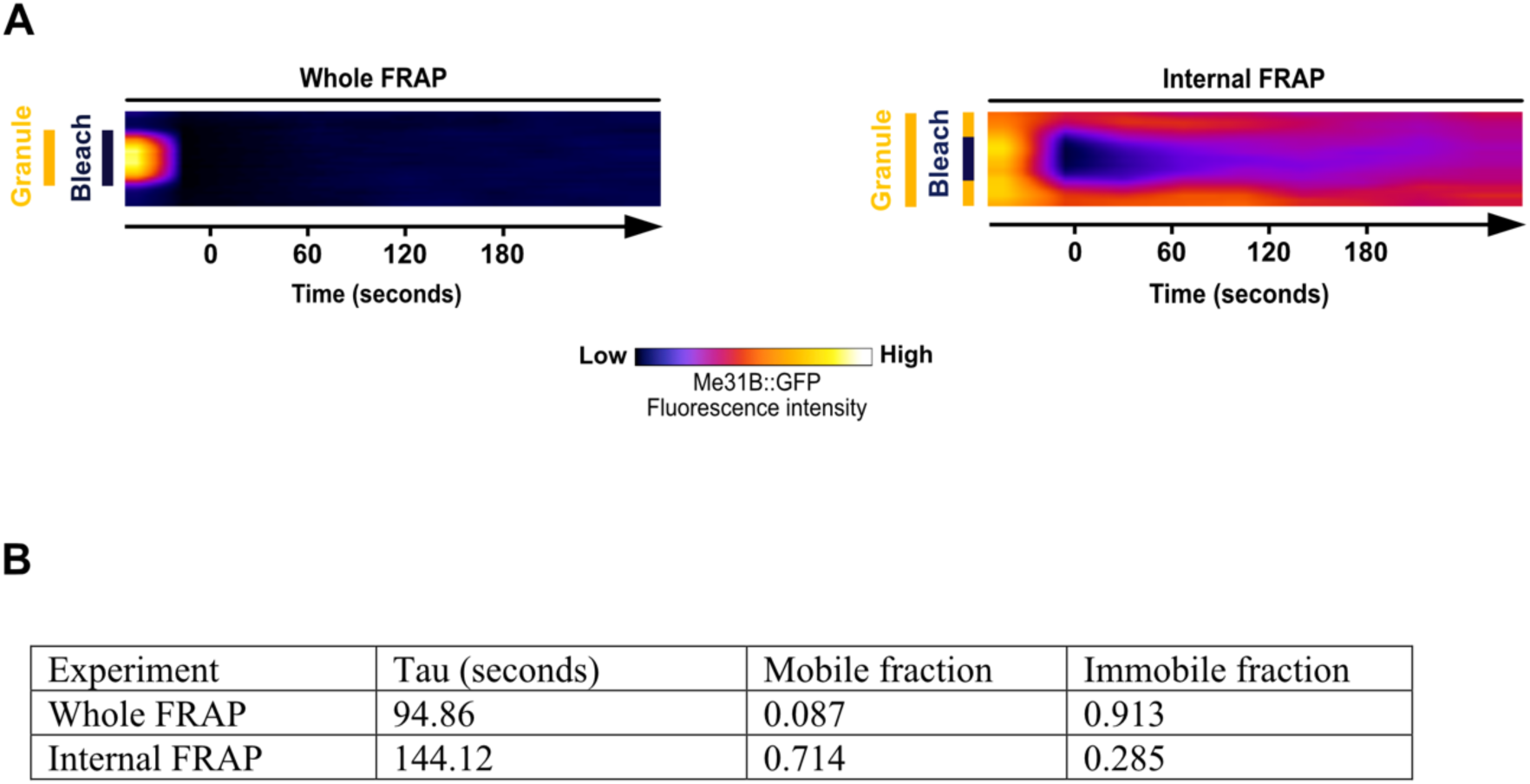
Me31B is mobile within P bodies but not between the cytoplasm and P bodies. **(A)** Kymograph of P body whole FRAP shows no recovery, while kymograph of P bodies post-internal FRAP displays recovery from the periphery to the center. This pattern is indicative of diffusion mediated recovery. **(B)** The tau, mobile and immobile fractions of Me31B after whole FRAP and internal FRAP experiment. These data show that Me31B is more mobile internally than between the P body and cytoplasm.

**Figure S3:**
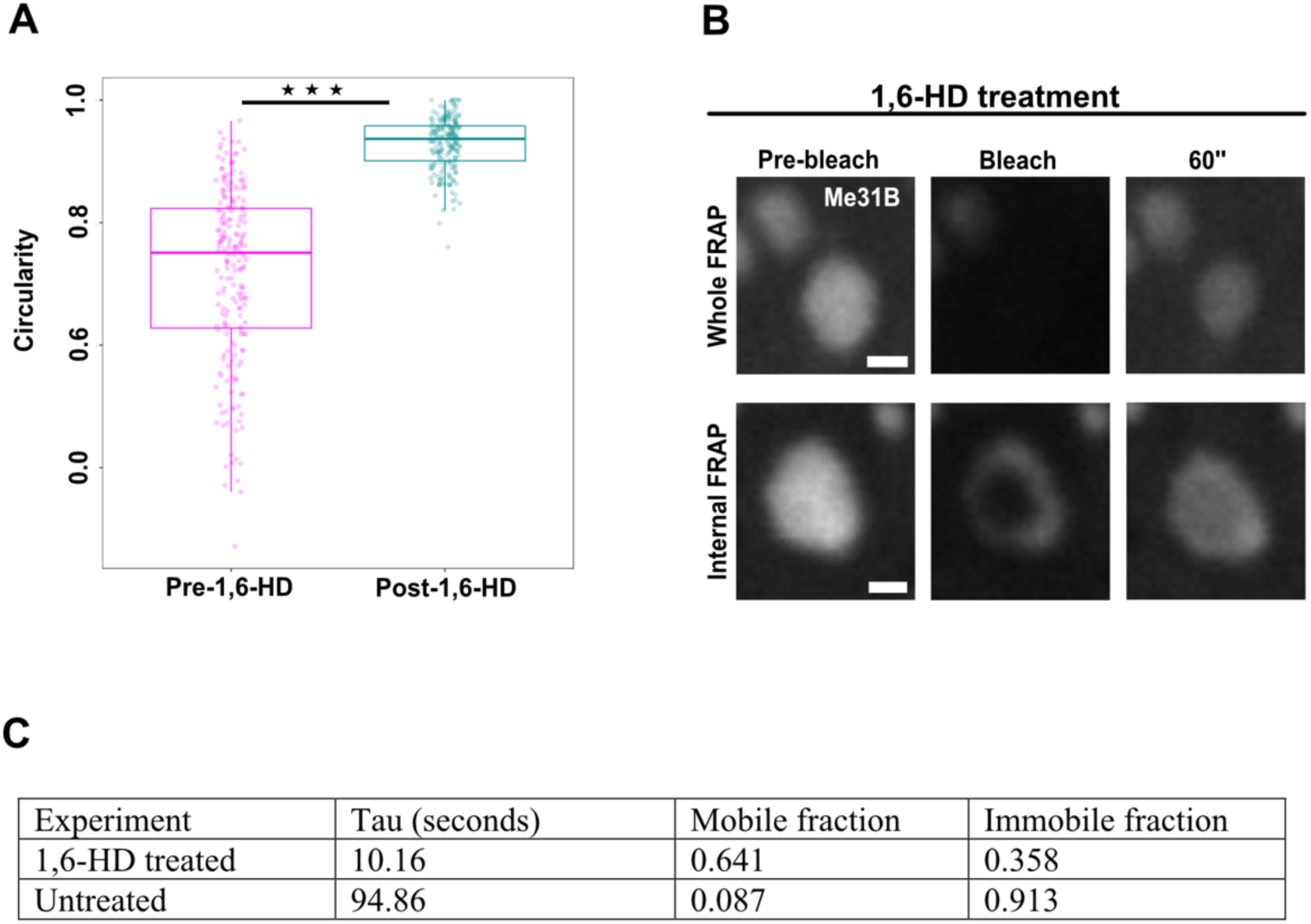
1,6-HD induces transition of P bodies to a more dynamic physical state. **(A)** Quantification of P body circularity before and after 1,6-HD treatment shows a significant increase in spherical morphology of P body post-treatment (Wilcoxon signed-rank test, n = 200). **(B)** Time series of P body condensates subjected to whole or internal FRAP post 1,6-HD treatment, both displaying rapid fluorescence recovery (n = 12 whole FRAP, n = 3 internal FRAP.) Scale bar: 3µm. **(C)** The tau, mobile and immobile fractions of Me31B after whole FRAP with or without the addition of 1,6-HD. These data show that Me31B is highly mobile between the P body and cytoplasm post-treatment.

**Figure S4:**
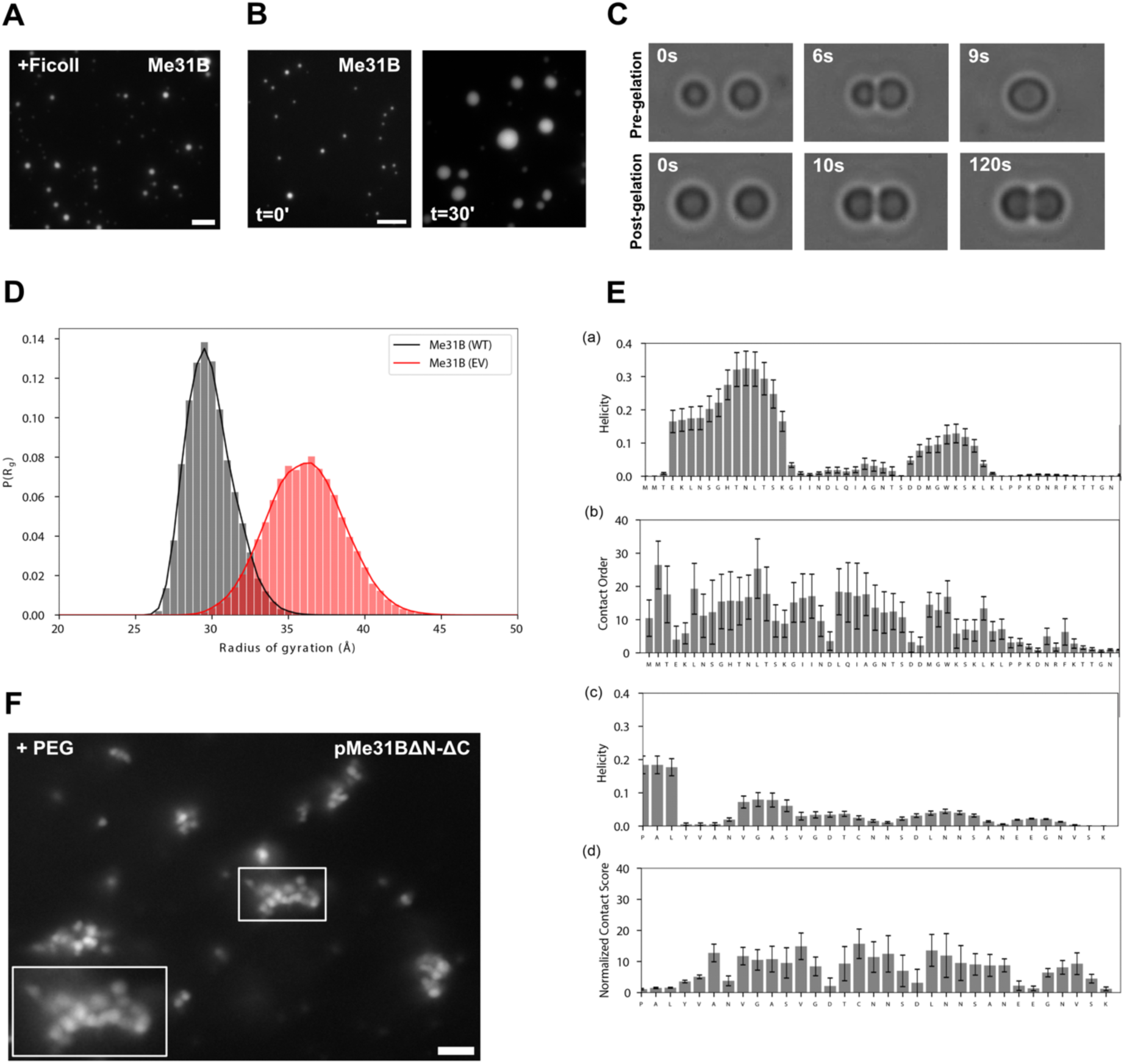
Interaction of IDRs with folded domains regulates Me31B physical state *in vitro*. **(A)** 7.5 µm purified GFP-pMe31B in the presence of 1% Ficoll readily forms phase separated condensates. **(B)** Time series of purified GFP-pMe31B (7.5 µm) shows increase in size of condensates over 30 minutes. **(C)** Time series of GFP-pMe31B condensate coalescence using optical traps. Rapid fusion of condensates is observed pre-gelation while condensates fail to fuse post-gelation. **(D)** Black bars show the Rg distribution for full-length Me31B under standard simulation conditions (full Hamiltonian (WT)), while red bars show the analogous distribution of the radius of gyration for simulations performed in which all attractive interactions are turned off (excluded volume (EV)). The full Hamiltonian simulations are substantially more compact, with an ensemble average radius of gyration of 30.1 Å, compared to 36.4 Å for the EV simulations. This compaction of the global dimensions originates from favorable interaction between the two IDRs and the folded domains. **(E)** Local helicity (a,c) and intramolecular contacts (b,d) quantified on a per-residue basis for the NTD (a,b) and CTD (c,d). The NTD possess two short transient helices (4-17 and 31-37), while the CTD is entirely devoid of secondary structures. The NTD engages in more extensive intramolecular interactions than the CTD, as quantified by the normalized contact score (d) (see methods, larger values mean more contacts per residue). In both cases, the IDRs engage relatively uniformly, as opposed to via a specific motif. This implies broad and non-specific interactions between the IDRs and the folded domains. **(F)** pMe31B with both IDRs deleted (pMe31B ΔN-ΔC), in the presence of 1% PEG, fails to coalesce and rather forms amorphous aggregates. Scale bar = 5µm (A,B,F), 1.5 µm (C).

**Figure S5:**
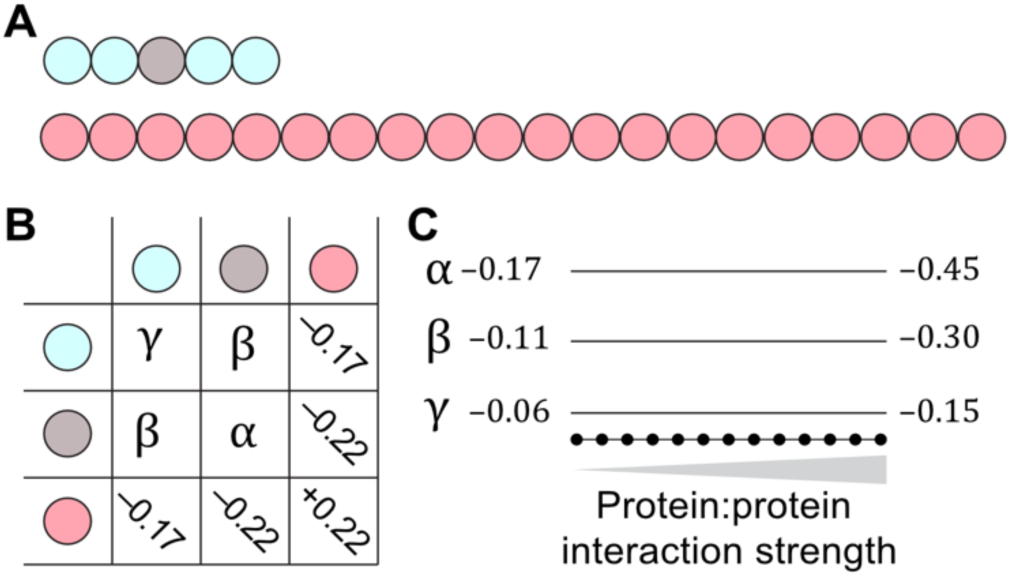
Schematic of course–grained simulations. **(A)** Topology of Me31b (top) and RNA (bottom) molecules used in coarse-grained simulations. **(B)** Basic interaction table showing relative bead-bead interaction strengths with protein:protein bead interactions defined in terms of parameters. **(C)** Protein:protein interaction parameters scale uniformly between max and min values.

## MATERIALS AND METHODS

### Drosophila stocks

The following transgenic lines were used in this paper:

Me31B::GFP (BL 51530, (Buszczak et al., 2007)), *hsp83-MCP-RFP* and *bcd-*(*ms2*)*_6_* (Weil et al., 2006), GFP::Tral (DGRC 110584, (Morin et al., 2001)), *tral^1^* (BDSC 14933) and *Df(3L)ED4483* (BDSC 8070) (Wilhelm et al., 2005).

Fly stocks were maintained at 25°C on Iberian recipe fly food as per standard procedure. For mature oocytes experiments, female flies were fed on yeast for two days at 25°C prior to dissection.

### Sample preparation

Mature oocytes from fattened female flies were dissected into halocarbon oil on a 22mm by 40mm cover slip for live imaging. For extrusion assays, membranes of dissected mature oocytes were poked and ruptured using sharp forceps to extrude the oocyte contents into the oil. Extruded material was then subjected to live imaging.

### Live imaging

Live imaging of *in vivo* and *ex vivo* P bodies, including all Fluorescent Recovery After Photobleaching experiments, were performed on the Olympus FV3000 microscope using the 1.35 NA, 60X silicone objective. Live imaging of recombinant Me31B condensates, induced on 35mm glass bottom MatTek dishes, was performed on the DeltaVision Core widefield microscope using a 1.4 NA, 60X oil immersion objective.

### Pharmacological treatments

Mature oocytes mounted in oil on a 22 by 40mm coverslip and set up under the microscope were treated with one or two drops of 10 µg/ml cytochalasin-D (Sigma-Aldrich) or 5% 1,6 hexanediol (Sigma-Aldrich) or 500ng/ml RNase A (Sigma-Aldrich) mixed in 1X Dulbecco’s Phosphate-buffered saline (PBS) solution without MgCl_2_ (Sigma-Aldrich), or home-made activation buffer (3.3mM NaH_2_PO_4_, 16.6mM KH_2_PO_4_, 10mM NaCl, 50mM KCl, 5% PEG 8000, 2mM CaCl_2_, pH 6.4; York-andersen *et al*., 2015) using a glass pipette. Me31B or Tral labelled P bodies before and after treatment were then imaged. For salt experiments, mature oocytes were extruded into various concentrations of MgCl_2_ or NaCl mixed in 1X PBS for 15 minutes before being subjected to live imaging. In the case of excessive movement of the oocytes or extruded material during addition of solutions, the focal plane of interest was adjusted accordingly, and imaging was performed.

### Protein purification

Recombinant GFP-Me31B, both the wild-type (WT) and the Me31BΔN-ΔC (mutant), were expressed in and purified from insect cells using the FlexiBAC baculovirus vector system (Lemaitre et al. 2019). Cell lysis was performed using a LM20 microfluidizer in lysis buffer containing 50mM Tris/HCl pH 7.6, 2mM EDTA, 1x EDTA-containing protease inhibitor cocktail (Roche), 1M KCl, 5% glycerol, 10mM imidazole, 3 ug/L Benzonase, 1 mM DTT. The soluble lysate fraction was collected after centrifugation for 1 hour at 16000 rpm (Beckman Coulter JA-25.50) at 4°C. MBP-tagged protein was captured by gravity flow affinity chromatography using amylose resin (New England Biolabs). Captured protein was washed with wash buffer (50mM Tris/HCl pH 7.6, 2mM EDTA,1M KCl, 5% glycerol, 10mM imidazole, 1 mM DTT, 3 ug/L Benzonase) and eluted using wash buffer containing 20mM maltose. The eluted protein was incubated with GST 3C-precision protease (1:50) at room temperature for 2 hours to cleave off affinity tags. Samples were applied to size exclusion chromatography using a HiLoad 16/600 Superdex 200 pg (GE Life Sciences) on an Akta pure chromatography system in 50mM Tris/HCl pH 7.6, 2mM EDTA,1M KCl, 5% glycerol, 1 mM DTT. Proteins were finally concentrated using an Amicon Ultra centrifugal-500-30K filter at 4000 xg. Aliquots were flash frozen and stored at −80°C.

### *In vitro* phase separation assay

Stored protein samples were thawed and spun to remove any residual precipitates. To induce phase separation of Me31B condensates (WT and mutant), 7.5 µM recombinant GFP-Me31B protein was added to an eppendorf tube containing the phase separation buffer(50mM KCl, 20mM PIPES, pH 7, 1% PEG-3K). Note: Gentle tapping of the tube induced phase separated, spherical condensates. Avoid mixing the content with a pipette tip as it induces aggregate formation.

### Optical tweezer experiments

Condensate fusions for wildtype or mutant condensates were quantified using a custom built dual-trap optical tweezer instrument (Jahnel et al., 2011). Condensates were induced in the phase separation buffer containing 5% PEG-3K at the following concentrations protein concentrations: For both WT and mutant condensates, 20 µM Me31B was used. Post condensation, two condensates were trapped using two separate optical traps and brought into close contact to induce fusion.

For quantifying the scaled fusion time for WT condensates, firstly, a relaxation time constant was derived from the fusion process over time. The scaled fusion time was then calculated by dividing the time constant by condensate radii to express the fusion time as a function independent of condensate size. For mutant condensates, due to their rapid aggregation post condensation, fusion was not quantifiable.

### Fluorescence Recovery After Photobleaching (FRAP)

For whole FRAP, Me31B/Tral labeled P bodies or *in vitro* Me31B condensates were entirely photobleached for 5 seconds using 40% laser intensity from the 405nm laser channel. For internal FRAP, a small region within Me31B labeled *in vivo* P bodies or *in vitro* Me31B condensates was photobleached for 5 seconds using 40% laser intensity from the 405nm laser channel. Time lapse series of Me31B fluorescence recovery was recorded every 30 seconds (*in vivo* P bodies) or 10 seconds (*in vitro* Me31B condensates) using the pre-bleach imaging parameters (minimal laser intensity using the 488nm laser channel, 2 Airy unit pinhole, 2048*2048 pixels). Mean fluorescence intensities were estimated using the Fiji ImageJ software. For whole FRAP analysis, background correction was performed by dividing Me31B fluorescent intensities of bleached condensates by fluorescent intensities of unbleached, cytoplasm. For internal FRAP, background correction was performed by dividing Me31B fluorescent intensities of bleached region within condensates by fluorescent intensities of whole condensates.

For all FRAP series, statistical analysis, curve fitting and plotting was performed using Rstudio/R software. Data for each condition was averaged and standard deviation was calculated where applicable. Recovery fitting of the normalized mean intensity as function of time was fitted by the least square analysis to determine fit to the single exponential equation: Normalized intensity = P×(1−e(−t/τ)) +y_0_ where y_0_ is the recovery plateau, t is time, τ is the time constant and P is the amplitude of the fluorescence change.

To infer the spatiotemporal pattern of fluorescence recovery, kymographs were produced using the ImageJ plugin ‘reslice’ by measuring fluorescence across of a region of interest over time.

### Fluorescence intensity measurements

Analysis of Me31B and *bcd* fluorescence before and after 1,6-HD treatment was performed using ImageJ processing software. Identical imaging parameters were utilized during imaging and measurement of fluorescence using ‘analyze particles’ and ‘measure’ feature on ImageJ. Individual Me31B and *bcd* particles were manually counted and analyzed before and after treatment with 1,6-HD.

### All-atom simulations

All-atom simulations were run with the ABSINTH implicit solvent model and the CAMPARI Monte Carlo simulation (V3.0) (http://campari.sourceforge.net/V3/index.html) and with the ion parameters derived by Mao *et al* (Mao and Pappu, 2012; Vitalis and Pappu, 2009). Preferential sampling is used such that the backbone dihedral angles of folded domains are held fixed, while all sidechain dihedral angles and the backbone dihedrals of folded proteins are fully sampled. In this way we *a priori* ensure that the folded domains remain folded. While the combination of ABSINTH and CAMPARI is well-established route to obtain reliable ensembles of disordered regions, more positional restraints on folded domains have been used previously applied to obtain good agreement with experiment (Cubuk et al., 2020; Martin and Mittag, 2018; Martin et al., 2020; Newcombe et al., 2018).

Starting structures were generated first by constructing homology models of Me31B based on the DDX6 structure (PDB: 4CT5) using SWISS-MODEL (Waterhouse et al., 2018). N- and C-terminal IDRs were constructed using CAMPARI. For all simulations, disordered regions were started from randomly generated non-overlapping random-coil conformations, with each replica using a unique starting structure. Monte Carlo simulations evolve the system via a series of moves that perturb backbone and sidechain dihedral angles along with the rigid-body coordinates of both polypeptides and explicit ions. Simulation analysis was performed using CAMPARITraj (www.ctraj.com) and MDTraj (McGibbon et al., 2015). The protein secondary structure was assessed using the DSSP algorithm (Kabsch and Sander, 1983).

Contact score analysis was performed by assessing the fraction of simulations in which two residues were in direct contact, a distance calibrated as 5.0 Å or shorter between heavy atoms. This fraction was divided by the analogous fraction computed from simulations in which all attractive molecular interactions (solvation effects, electrostatics, attractive component of the Lennard-Jones potential) were set to 0.0, in the so-called excluded volume (EV) limit (Holehouse et al., 2015).

All simulations were run at 10 mM NaCl and 310 K. Fifty independent simulations were run for a total of 80 million Monte Carlo steps with 5 million steps for equilibration. The system state saved every 100,000 steps. Each simulation generated 750 frames, generating a final ensemble of 37,500 frames. Where included, error bars are standard error of the mean over the fifty independent simulations.

### Bioinformatics

Disordered regions were calculated using both Mobidb-lite 1 and with metapredict 2,3. Disordered regions were identified using consensus scores from Mobidib-lite with a minimum IDR length of 25 residues and 3 or more predictors predicting a region to be disordered. The raw set of disordered regions from the drosophila proteome, along with analogous data for P body proteins is provided in the supplementary repository: https://github.com/holehouse-lab/supportingdata/tree/master/2021/sankaranarayanan_me31b_2021. Sequence analysis was performed using localCIDER4,5.

### Coarse-grained simulations

Coarse-grained simulations were performed with the PIMMS simulation engine. 6,7. Lattice-based Monte Carlo simulations afford a computationally tractable approach to sample systems with coexisting liquid phases, as has been applied in several different contexts 7–10. Monte Carlo moves include chain translate, rotate, and local/global pivot moves.

Simulations were run using a simple representation scheme in which Me31b was represented as a five-bead model made up of two N-terminal beads, a single central bead, and two C-terminal beads (Figure S4GA). In this way, the protein consists of intrinsically disordered region (IDR) beads and ordered domain (OD) beads. RNA is represented as a 20-bead homopolymer (Figure S4GA). We emphasize that these models are designed to describe a class of phenomenon, as opposed to capturing features specific to Me31b over RNA binding proteins. Our simplification of RNA and protein not-withstanding, these simple models allow us to interrogate general behavior.

The strength of interactions between the three bead types is shown in Figure S4GB and SXC. Units are in per kT (where k=1). The core keyfiles and parameter files used to run these simulations are provided at https://github.com/holehouse-lab/supportingdata/tree/master/2021/sankaranarayanan_me31b_2021.

For protein:RNA and RNA:RNA interaction strengths are held fixed across all simulations, while the protein:protein interaction strength is systematically altered across the simulations shown in Figure S4GC. The specific interaction strengths were chosen to qualitatively reflect insights from experimental work - i.e. OD:OD interaction is stronger than OD:IDR interaction, with IDR:IDR interaction being the weakest. We also assume both OD and IDR beads can interact with RNA, and that RNA:RNA interaction is repulsive. As a final note, we anticipate that RNA:RNA interactions plays an additional role in P body stability, assembly, and disassembly. However, for our initial simple model, absent of other specific information, we avoided adding more tunable parameters to develop a simple yet physically reasonable.

All simulations were run with 800 Me31b protein molecules. Simulations with RNA were also run with 50 RNA molecules. These numbers were chosen to ensure that reasonable statistics on droplet recruitment could be obtained with a sufficiently large system where bona fide phase separation occurs. Simulations were run on a 60 x 60 x 60 lattice with periodic boundary conditions, and simulation analysis was performed on the terminal 20% of the frames. Simulations were run for around 2.5 billion Monte Carlo moves, and three independent replicas were performed, such that error bars are the standard error of the mean on these replicas.

## ACKNOWLEDGEMENTS

We thank the Bloomington Drosophila Stock Center, Ilan Davis, Liz Gavis, James Wilhelm and William Chia for fly stocks. We are grateful to Paul Conduit for experimental advice; Titus Franzmann for advice on FRAP analysis, discussions and feedback on the manuscript; the Zoology Imaging Facility for assistance and support with microscopy; Protein Purification Facility at the MPI-CBG, Dresden; and funding from the University of Cambridge ISSF (097814) and the Wellcome Trust (200734/Z/16/Z) (to TTW), INLAKS and Cambridge Trust scholarship (to M.S.), and EMBO short-term fellowship, Sidney Sussex College, and Company of Biologists grants for travel and accommodation at the MPI-CBG, Dresden (to M.S.).

## AUTHOR CONTRIBUTIONS

M.S. performed the majority of experiments. M.S. and T.T.W. designed the majority of the experiments. R.J.E. and A.S.H. performed all simulations and *in silico* modeling. I.R.E.A.T. assisted with protein purification. M.J. performed the optical tweezer experiments. M.S., R.J.E., M.J., M.W., and A.S.H. analyzed the data. M.S. and T.T.W. wrote the manuscript. M.S., A.S.H., S.A. and T.T.W. edited the manuscript.

## REFERENCES

1. Andersen, A.H.Y., Wolfner, M.F., Wood, B.W., and Weil, T.T. (2020). A calcium - mediated actin redistribution at egg activation in Drosophila. 293–304.

2. Andrei, M.A., Ingelfinger, D., Heintzmann, R., Achsel, T., Rivera-pomar, R., and Lührmann, R. (2005). A role for eIF4E and eIF4E-transporter in targeting mRNPs to mammalian processing bodies. 717–727.

3. Bakthavachalu, B., Huelsmeier, J., Sudhakaran, I.P., Hillebrand, J., Singh, A., Petrauskas, A., Thiagarajan, D., Sankaranarayanan, M., Mizoue, L., Anderson, E.N., et al. (2018). RNP-Granule Assembly via Ataxin-2 Disordered Domains Is Required for Long-Term Memory and Neurodegeneration. Neuron 98, 754–766.e4.

4. Banani, S.F., Lee, H.O., Hyman, A.A., and Rosen, M.K. (2017). Biomolecular condensates: Organizers of cellular biochemistry. Nat. Rev. Mol. Cell Biol. 18, 285– 298.

5. Boeynaems, S., Alberti, S., Fawzi, N.L., Mittag, T., Polymenidou, M., Rousseau, F., Schymkowitz, J., Shorter, J., Wolozin, B., Van Den Bosch, L., et al. (2018). Protein Phase Separation: A New Phase in Cell Biology. Trends Cell Biol. 28, 420– 435.

6. Boeynaems, S., Holehouse, A.S., Weinhardt, V., Kovacs, D., Van Lindt, J., Larabell, C., Bosch, L. Van Den, Das, R., Tompa, P.S., Pappu, R. V., et al. (2019). Spontaneous driving forces give rise to protein−RNA condensates with coexisting phases and complex material properties. Proc. Natl. Acad. Sci. U. S. A. 116, 7889– 7898.

7. Boke, E., Ruer, M., Wühr, M., Coughlin, M., Lemaitre, R., Gygi, S.P., Alberti, S., Drechsel, D., Hyman, A.A., and Mitchison, T.J. (2016). Amyloid-like Self-Assembly of a Cellular Compartment. Cell 166, 637–650.

8. Bouveret, E. (2000). A Sm-like protein complex that participates in mRNA degradation. EMBO J. 19, 1661–1671.

9. Brangwynne, C.P., Eckmann, C.R., Courson, D.S., Rybarska, A., Hoege, C., Gharakhani, J., Jülicher, F., and Hyman, A.A. (2009). Germline P granules are liquid droplets that localize by controlled dissolution/condensation. Science (80-.). 324, 1729–1732.

10. Brangwynne, C.P., Mitchison, T.J., and Hyman, A.A. (2011). Active liquid-like behavior of nucleoli determines their size and shape in Xenopus laevis oocytes. Proc. Natl. Acad. Sci. U. S. A. 108, 4334–4339.

11. Brangwynne, C.P., Tompa, P., and Pappu, R. V. (2015). Polymer physics of intracellular phase transitions. Nat. Phys. 11, 899–904.

12. Buchan, J.R., Buchan, R., and Buchan, J.R. (2014). mRNP granules Assembly, function, and connections with disease. 6286.

13. Buszczak, M., Paterno, S., Lighthouse, D., Bachman, J., Planck, J., Owen, S., Skora, A.D., Nystul, T.G., Ohlstein, B., Allen, A., et al. (2007). The carnegie protein trap library: a versatile tool for Drosophila developmental studies. Genetics 175, 1505–1531.

14. Cubuk, J., Alston, J.J., Incicco, J.J., Singh, S., Stuchell-Brereton, M.D., Ward, M.D., Zimmerman, M.I., Vithani, N., Griffith, D., Wagoner, J.A., et al. (2020). The SARS-CoV-2 nucleocapsid protein is dynamic, disordered, and phase separates with RNA. BioRxiv.

15. Dorone, Y., Boeynaems, S., Jin, B., Bossi, F., Flores, E., and Lazarus, E. (2020). Hydration-dependent phase separation of a prion-like protein regulates seed germination during water stress.

16. Dzuricky, M., Rogers, B.A., Shahid, A., Cremer, P.S., and Chilkoti, A. (2020). using artificial disordered proteins. Nat. Chem. 12.

17. Eichhorn, S.W., Subtelny, A.O., Kronja, I., Kwasnieski, J.C., Orr-Weaver, T.L., and Bartel, D.P. (2016). mRNA poly(A)-tail changes specified by deadenylation broadly reshape translation in Drosophila oocytes and early embryos. Elife 5, 1–24.

18. Eulalio, A., Behm-Ansmant, I., Schweizer, D., and Izaurralde, E. (2007). P-Body Formation Is a Consequence, Not the Cause, of RNA-Mediated Gene Silencing. Mol. Cell. Biol. 27, 3970–3981.

19. Feric, M., Vaidya, N., Harmon, T.S., Mitrea, D.M., Zhu, L., Richardson, T.M., Kriwacki, R.W., Pappu, R. V., and Brangwynne, C.P. (2016). Coexisting Liquid Phases Underlie Nucleolar Subcompartments. Cell 165, 1686–1697.

20. Franzmann, T.M., Jahnel, M., Pozniakovsky, A., Mahamid, J., Holehouse, A.S., Nüske, E., Richter, D., Baumeister, W., Grill, S.W., Pappu, R. V., et al. (2018). Phase separation of a yeast prion protein promotes cellular fitness. Science (80). 359.

21. Götze, M., Dufourt, J., Ihling, C., Rammelt, C., Pierson, S., Sambrani, N., Temme, C., Sinz, A., Simonelig, M., and Wahle, E. (2017). Translational repression of the Drosophila nanos mRNA involves the RNA helicase Belle and RNA coating by Me31B and Trailer hitch. 1552–1568.

22. Hara, M., Lourido, S., Petrova, B., Lou, H.J., Von Stetina, J.R., Kashevsky, H., Turk, B.E., and Orr-Weaver, T.L. (2018). Identification of PNG kinase substrates uncovers interactions with the translational repressor TRAL in the oocyte-to-embryo transition. Elife 7, 1–19.

23. Holehouse, A.S., Garai, K., Lyle, N., Vitalis, A., and Pappu, R. V (2015). Quantitative assessments of the distinct contributions of polypeptide backbone amides versus side chain groups to chain expansion via chemical denaturation. J. Am. Chem. Soc. 137, 2984–2995.

24. Hondele, M., Sachdev, R., Heinrich, S., Wang, J., Vallotton, P., Fontoura, B.M.A., and Weis, K. (2019). DEAD-box ATPases are global regulators of phase-separated organelles. Nature 573, 144–148.

25. Hubstenberger, A., Noble, S.L., Cameron, C., and Evans, T.C. (2013). Translation repressors, an RNA helicase, and developmental cues control RNP phase transitions during early development. Dev. Cell 27, 161–173.

26. Hubstenberger, A., Courel, M., Bénard, M., Souquere, S., Ernoult-Lange, M., Chouaib, R., Yi, Z., Morlot, J.B., Munier, A., Fradet, M., et al. (2017). P-Body Purification Reveals the Condensation of Repressed mRNA Regulons. Mol. Cell 68, 144–157.e5.

27. Hyman, A.A., Weber, C.A., and Jülicher, F. (2014). Liquid-Liquid Phase Separation in Biology. Annu. Rev. Cell Dev. Biol. 30, 39–58.

28. Jahnel, M., Behrndt, M., Jannasch, A., Schäffer, E., and Grill, S.W. (2011). Measuring the complete force field of an optical trap. Opt. Lett. 36, 1260–1262.

29. Jung, H., Gkogkas, C.G., Sonenberg, N., and Holt, C.E. (2014). Review Remote Control of Gene Function by Local Translation. Cell 157, 26–40.

30. Kabsch, W., and Sander, C. (1983). Dictionary of protein secondary structure: pattern recognition of hydrogen-bonded and geometrical features. Biopolymers 22, 2577–2637.

31. Kaneuchi, T., Sartain, C. V, Takeo, S., Horner, V.L., Buehner, N.A., and Aigaki, T. (2015). Calcium waves occur as Drosophila oocytes activate. 112, 791–796.

32. Kato, M., Han, T.W., Xie, S., Shi, K., Du, X., Wu, L.C., Mirzaei, H., Goldsmith, E.J., Longgood, J., Pei, J., et al. (2012). Cell-free formation of RNA granules: Low complexity sequence domains form dynamic fibers within hydrogels. Cell 149, 753–767.

33. Kedersha, N., Stoecklin, G., Ayodele, M., Yacono, P., Lykke-andersen, J., Marvin, J.F., Scheuner, D., Kaufman, R.J., Golan, D.E., and Anderson, P. (2005). Stress granules and processing bodies are dynamically linked sites of mRNP remodeling 4. 169, 871–884.

34. Kloc, M., and Etkin, L.D. (2005). RNA localization mechanisms in oocytes. J. Cell Sci. 118, 269–282.

35. Krauchunas, A.R., and Wolfner, M.F. (2013). Molecular Changes During Egg Activation. Curr. Top. Dev. Biol. 102, 267–292.

36. Kroschwald, S., Maharana, S., Mateju, D., Malinovska, L., Nüske, E., Poser, I., Richter, D., and Alberti, S. (2015). Promiscuous interactions and protein disaggregases determine the material state of stress-inducible RNP granules. Elife 4, 1–32.

37. Kroschwald, S., Munder, M.C., Maharana, S., Ruer, M., Hyman, A.A., Alberti, S., Kroschwald, S., Munder, M.C., Maharana, S., Franzmann, T.M., et al. (2018). Different Material States of Pub1 Condensates Define Distinct Modes of Stress Adaptation and Article Different Material States of Pub1 Condensates Define Distinct Modes of Stress Adaptation and Recovery. CellReports 23, 3327–3339.

38. Lasko, P. (2012). mRNA localization and translational control in Drosophila oogenesis. Cold Spring Harb. Perspect. Biol. 4, 1–15.

39. Li, P., Banjade, S., Cheng, H.C., Kim, S., Chen, B., Guo, L., Llaguno, M., Hollingsworth, J. V., King, D.S., Banani, S.F., et al. (2012). Phase transitions in the assembly of multivalent signalling proteins. Nature 483, 336–340.

40. Lin, M. Der, Jiao, X., Grima, D., Newbury, S.F., Kiledjian, M., and Chou, T. Bin (2008). Drosophila processing bodies in oogenesis. Dev. Biol. 322, 276–288.

41. Lin, Y., Protter, D.S.W., Rosen, M.K., and Parker, R. (2015a). Formation and Maturation of Phase-Separated Liquid Droplets by RNA-Binding Proteins. Mol. Cell 60, 208–219.

42. Lin, Y., Protter, D.S.W., Rosen, M.K., and Parker, R. (2015b). Formation and Maturation of Phase-Separated Liquid Droplets by RNA-Binding Proteins. Mol. Cell 60, 208–219.

43. Luo, Y., Na, Z., and Slavoff, S.A. (2018). P-Bodies: Composition, Properties, and Functions. Biochemistry 57, 2424–2431.

44. Lyon, A.S., Peeples, W.B., and Rosen, M.K. (2021). P H A S E S E PA R AT I O N I N B I O LO G Y. Nat. Rev. Mol. Cell Biol. 22.

45. Majumdar, A., Cesario, W.C., White-Grindley, E., Jiang, H., Ren, F., Khan, M. “Repon,” Li, L., Choi, E.M.-L., Kannan, K., Guo, F., et al. (2012). Critical Role of Amyloid-like Oligomers of Drosophila Orb2 in the Persistence of Memory. Cell 148, 515–529.

46. Mao, A.H., and Pappu, R. V (2012). Crystal lattice properties fully determine short-range interaction parameters for alkali and halide ions. J. Chem. Phys. 137, 64104.

47. Martin, E.W., and Mittag, T. (2018). Relationship of Sequence and Phase Separation in Protein Low-Complexity Regions. Biochemistry 57, 2478–2487.

48. Martin, E.W., Holehouse, A.S., Peran, I., Farag, M., Incicco, J.J., Bremer, A., Grace, C.R., Soranno, A., Pappu, R. V., and Mittag, T. (2020). Valence and patterning of aromatic residues determine the phase behavior of prion-like domains. Science (80-.). 367, 694–699.

49. McCambridge, A., Solanki, D., Olchawa, N., Govani, N., Trinidad, J.C., and Gao, M. (2020). Comparative Proteomics Reveal Me31B’s Interactome Dynamics, Expression Regulation, and Assembly Mechanism into Germ Granules during Drosophila Germline Development. Sci. Rep. 10, 1–13.

50. McGibbon, R.T., Beauchamp, K.A., Harrigan, M.P., Klein, C., Swails, J.M., Hernández, C.X., Schwantes, C.R., Wang, L.-P., Lane, T.J., and Pande, V.S. (2015). MDTraj: A Modern Open Library for the Analysis of Molecular Dynamics Trajectories. Biophys. J. 109, 1528–1532.

51. Medioni, C., Mowry, K., and Besse, F. (2012). Principles and roles of mRNA localization in animal development. Dev. 139, 3263–3276.

52. Mitrea, D.M., Cika, J.A., Guy, C.S., Ban, D., Banerjee, P.R., Stanley, C.B., Nourse, A., Deniz, A.A., and Kriwacki, R.W. (2016). Nucleophosmin integrates within the nucleolus via multi-modal interactions with proteins displaying R-rich linear motifs and rRNA. 1–33.

53. Monzo, K., Papoulas, O., Cantin, G.T., Wang, Y., Yates, J.R., and Sisson, J.C. (2006). Fragile X mental retardation protein controls trailer hitch expression and cleavage furrow formation in Drosophila embryos. Proc. Natl. Acad. Sci. U. S. A. 103, 18160–18165.

54. Morin, X., Daneman, R., Zavortink, M., and Chia, W. (2001). A protein trap strategy to detect GFP-tagged proteins expressed from their endogenous loci in Drosophila. Proc. Natl. Acad. Sci. U. S. A. 98, 15050–15055.

55. Murthy, A.C., Dignon, G.L., Kan, Y., Zerze, G.H., Parekh, S.H., Mittal, J., and Fawzi, N.L. (2019). Molecular interactions underlying liquid − liquid phase separation of the FUS low-complexity domain. Nat. Struct. Mol. Biol. 26.

56. Nakamura, A., Amikura, R., Hanyu, K., and Kobayashi, S. (2001). Me31B silences translation of oocyte-localizing RNAs through the formation of cytoplasmic RNP complex during Drosophila oogenesis. Development 128, 3233–3242.

57. Newcombe, E.A., Ruff, K.M., Sethi, A., Ormsby, A.R., Ramdzan, Y.M., Fox, A., Purcell, A.W., Gooley, P.R., Pappu, R. V, and Hatters, D.M. (2018). Tadpole-like Conformations of Huntingtin Exon 1 Are Characterized by Conformational Heterogeneity that Persists regardless of Polyglutamine Length. J. Mol. Biol. 430, 1442–1458.

58. Nott, T.J., Petsalaki, E., Farber, P., Jervis, D., Fussner, E., Plochowietz, A., Craggs, T.D., Bazett-Jones, D.P., Pawson, T., Forman-Kay, J.D., et al. (2015). Phase Transition of a Disordered Nuage Protein Generates Environmentally Responsive Membraneless Organelles. Mol. Cell 57, 936–947.

59. Pak, C.W., Kosno, M., Holehouse, A.S., Padrick, S.B., Mittal, A., Ali, R., Yunus, A.A., Liu, D.R., Pappu, R. V, and Rosen, M.K. (2016). Sequence Determinants of Intracellular Phase Separation by Complex Coacervation of a Disordered Protein. Mol. Cell 63, 72–85.

60. Parker, R., and Sheth, U. (2007). P Bodies and the Control of mRNA Translation and Degradation. Mol. Cell 25, 635–646.

61. Patel, A., Lee, H.O., Jawerth, L., Maharana, S., Jahnel, M., Hein, M.Y., Stoynov, S., Mahamid, J., Saha, S., Franzmann, T.M., et al. (2015). A Liquid-to-Solid Phase Transition of the ALS Protein FUS Accelerated by Disease Mutation. Cell 162, 1066–1077.

62. Patel, S.S., Belmont, B.J., Sante, J.M., and Rexach, M.F. (2007). Natively Unfolded Nucleoporins Gate Protein Diffusion across the Nuclear Pore Complex. 83–96.

63. Powers, S.K., Holehouse, A.S., Korasick, D.A., Schreiber, K.H., Clark, N.M., Jing, H., Emenecker, R., Han, S., Tycksen, E., Hwang, I., et al. (2019). Nucleo-cytoplasmic Partitioning of ARF Proteins Controls Auxin Responses in Arabidopsis thaliana. Mol. Cell 76, 177–190.e5.

64. Protter, D.S.W., Rao, B.S., Treeck], B. [Van, Lin, Y., Mizoue, L., Rosen, M.K., and Parker, R. (2018). Intrinsically Disordered Regions Can Contribute Promiscuous Interactions to RNP Granule Assembly. Cell Rep. 22, 1401–1412.

65. Puthanveettil, S. V (2013). RNA transport and long-term memory storage. 6286.

66. Rajasethupathy, P., Fiumara, F., Sheridan, R., Betel, D., Puthanveettil, S. V, Russo, J.J., Sander, C., Tuschl, T., and Kandel, E. (2009). Article Characterization of Small RNAs in Aplysia Reveals a Role for miR-124 in Constraining Synaptic Plasticity through CREB. Neuron 63, 803–817.

67. Riback, J.A., Katanski, C.D., Kear-Scott, J.L., Pilipenko, E. V, Rojek, A.E., Sosnick, T.R., and Drummond, D.A. (2017). Stress-Triggered Phase Separation Is an Adaptive, Evolutionarily Tuned Response. Cell 168, 1028–1040.e19.

68. Ribbeck, K., and Go, D. (2002). The permeability barrier of nuclear pore complexes appears to operate via hydrophobic exclusion. 21.

69. Rog, O., Köhler, S., and Dernburg, A.F. (2017). The synaptonemal complex has liquid crystalline properties and spatially regulates meiotic recombination factors. Elife 6.

70. Shin, Y., and Brangwynne, C.P. (2017). Liquid phase condensation in cell physiology and disease. Science (80-.). 357.

71. Sudhakaran, I.P., and Ramaswami, M. (2017). Long-term memory consolidation: The role of RNA-binding proteins with prion-like domains. RNA Biol. 14, 568–586.

72. Tadros, W., and Lipshitz, H.D. (2009). The maternal-to-zygotic transition: a play in two acts. 3042, 3033–3042.

73. Tritschler, F., Eulalio, A., Truffault, V., Hartmann, M.D., Helms, S., Schmidt, S., Coles, M., Izaurralde, E., and Weichenrieder, O. (2007). A Divergent Sm Fold in EDC3 Proteins Mediates DCP1 Binding and P-Body Targeting. Mol. Cell. Biol. 27, 8600–8611.

74. Tritschler, F., Eulalio, A., Helms, S., Schmidt, S., Coles, M., Weichenrieder, O., Izaurralde, E., and Truffault, V. (2008). Similar Modes of Interaction Enable Trailer Hitch and EDC3 To Associate with DCP1 and Me31B in Distinct Protein Complexes. Mol. Cell. Biol. 28, 6695–6708.

75. Tritschler, F., Braun, J.E., Eulalio, A., Truffault, V., Izaurralde, E., and Weichenrieder, O. (2009). Structural Basis for the Mutually Exclusive Anchoring of P Body Components EDC3 and Tral to the DEAD Box Protein DDX6/Me31B. Mol. Cell 33, 661–668.

76. Vitalis, A., and Pappu, R. V (2009). ABSINTH: a new continuum solvation model for simulations of polypeptides in aqueous solutions. J. Comput. Chem. 30, 673– 699.

77. Wang, J.T., Smith, J., Chen, B., Schmidt, H., Rasoloson, D., Paix, A., Lambrus, B.G., Calidas, D., Betzig, E., and Seydoux, G. (2014). Regulation of RNA granule dynamics by phosphorylation of serine-rich, intrinsically disordered proteins in C. elegans. 1–23.

78. Wang, M., Ly, M., Lugowski, A., Laver, J.D., Lipshitz, H.D., Smibert, C.A., and Rissland, O.S. (2017). ME31B globally represses maternal mRNAs by two distinct mechanisms during the Drosophila maternal-to-zygotic transition. Elife 6, 1–22.

79. Waterhouse, A., Bertoni, M., Bienert, S., Studer, G., Tauriello, G., Gumienny, R., Heer, F.T., de Beer, T.A.P., Rempfer, C., Bordoli, L., et al. (2018). SWISS-MODEL: homology modelling of protein structures and complexes. Nucleic Acids Res. 46, W296–W303.

80. Weber, S.C. (2017). ScienceDirect Sequence-encoded material properties dictate the structure and function of nuclear bodies. Curr. Opin. Cell Biol. 46, 62–71.

81. Weber, S.C., and Brangwynne, C.P. (2012). Minireview Getting RNA and Protein in Phase. Cell 149, 1188–1191.

82. Weil, T.T., Forrest, K.M., and Gavis, E.R. (2006). Localization of bicoid mRNA in late oocytes is maintained by continual active transport. Dev. Cell 11, 251–262.

83. Weil, T.T., Parton, R.M., Herpers, B., Soetaert, J., Veenendaal, T., Xanthakis, D., Dobbie, I.M., Halstead, J.M., Hayashi, R., Rabouille, C., et al. (2012). Drosophila patterning is established by differential association of mRNAs with P bodies. Nat. Cell Biol. 14, 1305–1313.

84. Wilhelm, J.E., Buszczak, M., and Sayles, S. (2005). Efficient protein trafficking requires trailer hitch, a component of a ribonucleoprotein complex localized to the ER in Drosophila. Dev. Cell 9, 675–685.

85. Woodruff, J.B., Ferreira Gomes, B., Widlund, P.O., Mahamid, J., Honigmann, A., and Hyman, A.A. (2017). The Centrosome Is a Selective Condensate that Nucleates Microtubules by Concentrating Tubulin. Cell 169, 1066–1077.e10.

86. York-andersen, A.H., Parton, R.M., Bi, C.J., Bromley, C.L., Davis, I., and Weil, T.T. (2015). RESEARCH ARTICLE A single and rapid calcium wave at egg activation in Drosophila. 553–560.

87. Zhang, H., Elbaum-Garfinkle, S., Langdon, E.M., Taylor, N., Occhipinti, P., Bridges, A.A., Brangwynne, C.P., and Gladfelter, A.S. (2015). RNA Controls PolyQ Protein Phase Transitions. Mol. Cell 60, 220–230.

## MATERIALS AND METHODS REFERENCES

1. Buszczak, M., Paterno, S., Lighthouse, D., Bachman, J., Planck, J., Owen, S., Skora, A.D., Nystul, T.G., Ohlstein, B., Allen, A., et al. (2007). The carnegie protein trap library: a versatile tool for Drosophila developmental studies. Genetics 175, 1505–1531.

2. Cubuk, J., Alston, J.J., Incicco, J.J., Singh, S., Stuchell-Brereton, M.D., Ward, M.D., Zimmerman, M.I., Vithani, N., Griffith, D., Wagoner, J.A., et al. (2020). The SARS-CoV-2 nucleocapsid protein is dynamic, disordered, and phase separates with RNA. BioRxiv.

3. Holehouse, A.S., Garai, K., Lyle, N., Vitalis, A., and Pappu, R. V (2015). Quantitative assessments of the distinct contributions of polypeptide backbone amides versus side chain groups to chain expansion via chemical denaturation. J. Am. Chem. Soc. 137, 2984–2995.

4. Jahnel, M., Behrndt, M., Jannasch, A., Schäffer, E., and Grill, S.W. (2011). Measuring the complete force field of an optical trap. Opt. Lett. 36, 1260–1262.

5. Kabsch, W., and Sander, C. (1983). Dictionary of protein secondary structure: pattern recognition of hydrogen-bonded and geometrical features. Biopolymers 22, 2577–2637.

6. Mao, A.H., and Pappu, R. V (2012). Crystal lattice properties fully determine short-range interaction parameters for alkali and halide ions. J. Chem. Phys. 137, 64104.

7. Martin, E.W., and Mittag, T. (2018). Relationship of Sequence and Phase Separation in Protein Low-Complexity Regions. Biochemistry 57, 2478–2487.

8. Martin, E.W., Holehouse, A.S., Peran, I., Farag, M., Incicco, J.J., Bremer, A., Grace, C.R., Soranno, A., Pappu, R. V., and Mittag, T. (2020). Valence and patterning of aromatic residues determine the phase behavior of prion-like domains. Science (80). 367, 694–699.

9. McGibbon, R.T., Beauchamp, K.A., Harrigan, M.P., Klein, C., Swails, J.M., Hernández, C.X., Schwantes, C.R., Wang, L.-P., Lane, T.J., and Pande, V.S. (2015). MDTraj: A Modern Open Library for the Analysis of Molecular Dynamics Trajectories. Biophys. J. 109, 1528–1532.

10. Morin, X., Daneman, R., Zavortink, M., and Chia, W. (2001). A protein trap strategy to detect GFP-tagged proteins expressed from their endogenous loci in Drosophila. Proc. Natl. Acad. Sci. U. S. A. 98, 15050–15055.

11. Newcombe, E.A., Ruff, K.M., Sethi, A., Ormsby, A.R., Ramdzan, Y.M., Fox, A., Purcell, A.W., Gooley, P.R., Pappu, R. V, and Hatters, D.M. (2018). Tadpole-like Conformations of Huntingtin Exon 1 Are Characterized by Conformational Heterogeneity that Persists regardless of Polyglutamine Length. J. Mol. Biol. 430, 1442–1458.

12. Vitalis, A., and Pappu, R. V (2009). ABSINTH: a new continuum solvation model for simulations of polypeptides in aqueous solutions. J. Comput. Chem. 30, 673– 699.

13. Waterhouse, A., Bertoni, M., Bienert, S., Studer, G., Tauriello, G., Gumienny, R., Heer, F.T., de Beer, T.A.P., Rempfer, C., Bordoli, L., et al. (2018). SWISS-MODEL: homology modelling of protein structures and complexes. Nucleic Acids Res. 46, W296–W303.

14. Weil, T.T., Forrest, K.M., and Gavis, E.R. (2006). Localization of bicoid mRNA in late oocytes is maintained by continual active transport. Dev. Cell 11, 251–262.

15. Wilhelm, J.E., Buszczak, M., and Sayles, S. (2005). Efficient protein trafficking requires trailer hitch, a component of a ribonucleoprotein complex localized to the ER in Drosophila. Dev. Cell 9, 675–685.

16. York-andersen, A.H., Parton, R.M., Bi, C.J., Bromley, C.L., Davis, I., and Weil, T.T. (2015). A single and rapid calcium wave at egg activation in Drosophila. Biology Open 553–560.

